# Differences in berry primary and secondary metabolisms identified by transcriptomic and metabolic profiling of two table grape color somatic variants

**DOI:** 10.1101/861120

**Authors:** Claudia Santibáñez, Carlos Meyer, Litsy Martínez, Tomás Moyano, John Lunn, Regina Feil, Zhanwu Dai, David Carrasco, Rosa Arroyo-García, Ghislaine Hilbert, Christel Renaud, Serge Delrot, Fabiane Manke-Nachtigall, Rodrigo Gutiérrez, José Tomás Matus, Eric Gomès, Patricio Arce-Johnson

## Abstract

Anthocyanins are flavonoids responsible for the color of berries in skin-pigmented grapevine (*Vitis vinifera* L.). Due to the widely adopted vegetative propagation of this species, somatic mutations occurring in meristematic cell layers can be fixed and passed into the rest of the plant when cloned. In this study we focused on the transcriptomic and metabolic differences between two color somatic variants. Using microscopic, metabolic and mRNA profiling analyses we compared the table grape cultivar (cv.) ‘Red Globe’ (RG, with purplish berry skin) and cv. ‘Chimenti Globe’ (CG, with a contrasting reddish berry skin color). As expected, significant differences were found in the composition of flavonoids and other phenylpropanoids, but also in their upstream precursors’ shikimate and phenylalanine. Among primary metabolites, sugar phosphates related with sucrose biosynthesis were less accumulated in cv. ‘CG’. The red-skinned cv. ‘CG’ only contained di-hydroxylated anthocyanins (i.e. peonidin and cyanidin) while the tri-hydroxylated derivatives malvidin, delphinidin and petunidin were absent, in correlation to the reddish cv. ‘CG’ skin coloration. Transcriptomic analysis showed alteration in flavonoid metabolism and terpenoid pathways and in primary metabolism such as sugar content. Eleven *flavonoid 3’5’-hydroxylase* gene copies were down-regulated in cv. ‘CG’. This family of cytochrome P450 oxidoreductases are key in the biosynthesis of tri-hydroxylated anthocyanins. Many transcription factors appeared down-regulated in cv. ‘CG’ in correlation to the metabolic and transcriptomic changes observed. The use of molecular markers and its confirmation with our RNA-seq data showed the exclusive presence of the null *MYBA2* white allele (i.e. homozygous in both L1 and L2 layers) in the two somatic variants. Therefore, the differences in *MYBA1* expression seem sufficient for the skin pigmentation differences and the changes in MYBA target gene expression in cv. ‘Chimenti Globe’.

## INTRODUCTION

Table grapes are produced for their fresh and dried (i.e. raisins) consumption, with their markets continuously demanding better fruit quality but also searching for the generation of new cultivars with appealing features. The berry developmental process displays a double sigmoid curve with two growth phases separated by a lag phase, at the end of which veraison takes place (COOMBE, 1995). The accumulation of pigments (i.e. secondary metabolites known as anthocyanins) occurs at this stage in red and black-skinned cultivars, concomitant with sugars (primary metabolites) and other secondary metabolites (e.g. aroma volatile compounds). All these accumulate until the fruit exhibits its desirable quality traits and it is ready for harvest.

Berry skin color is an important trait for the grapevine industry, used as a criterion for selection within breeding programs. Berry skin colors ranges from black to blue, red or pink and also yellowish-green tones and hues, as a consequence of natural hybridization and human selection processes (Azuma, 2018; Azuma et al., 2008; Frédérique Pelsy, Dumas, Bévilacqua, Hocquigny, & Merdinoglu, 2015). Clonal polymorphism affecting berry color is a common event that occurs in grapevines thanks to vegetative propagation. Under these circumstances somatic mutations that occur in the meristematic cells within buds (in the entire meristem -bud sports- or only a portion -chimeras-) are maintained and propagated, leading to somatic color variants (Pelsy et al., 2015; D’Amato, 1997). In most cases, somatic mutations affect only one cell layer of the meristem, leading to periclinal chimeras. This phenomenon engenders a heritable variation source creating new grape cultivars.

The main somatic polymorphisms accounting for color variation include insertions, duplications or SNPs, some caused by the activity of transposable elements disturbing regulatory regions (Carrier et al., 2012). Some well-known examples of clonal polymorphism affecting berry color correspond to the case of cultivars (cv.) ‘Pinot Noir’ and ‘Pinot Blanc’, with the later presenting a large deletion (over 260 kb-long) that removes the functional *MYBA1* and *MYBA2* genes that are the main regulators of anthocyanin synthesis (Walker et al., 2006; Yakushiji et al., 2006). A different somatic mutation was found in cv. ‘Pinot Gris’ but which affected the same two genes, producing a grey-skinned phenotype. In this second variant a chimeric structure was identified, with berries composed of a colored L1-derived epidermis (heterozygous for the functional and null alleles) while the L2 cells possessed a homozygous mutation in both *MYBA1* and *MYBA2* (Vezzulli et al., 2012).

The *MYBA1* and *MYBA2* genes form part of the berry color locus found in chromosome 2 (Hocquigny et al., 2004). These factors are known to form ternary complexes with bHLH (basic helix–loop–helix) and WDR (tryptophan–aspartic acid repeat) proteins (Hichri et al., 2010; Matus et al., 2010) being all of them important for the activation of the structural genes of the flavonoid branch in the phenylpropanoid pathway. Additional factors have been identified to date in this species (Matus et al., 2017), demonstrating a complex hierarchical regulation of pigment accumulation according to organ and in response to the environment.

In the last decade several genome-wide studies have demonstrated that a large transcriptomic shift drives the transition from the green berry to the ripening stages (Fasoli et al., 2012; Massonnet et al., 2017; Palumbo et al., 2014). The availability of a high quality grapevine genomic sequence and of large amount of transcriptomics and metabolomics data has allowed to conduct integrative analyses to decipher the key transcriptomic reprogramming events occurring at the onset and very late stages of berry ripening (reviewed by (Serrano et al., 2017; Wong & Matus, 2017). However, most of the transcriptomic comparisons conducted so far have considered very distant cultivars (Ghan et al., 2015; Massonnet et al., 2017; Zenoni et al., 2016) while very few have compared closely related cultivars at the genome-wide level (Carbonell-Bejerano et al., 2017). Here, we performed a microscopic, metabolic and transcriptomic characterization of berry development and ripening in two almost isogenic cultivars with different berry skin pigmentation. Berries from the cultivar ‘Red Globe’ (RG), characterized by a large size, thick skin and dark purple skin color, were compared to those of its somatic variant cv. ‘Chimenti Globe’ (CG) with similar characteristics but differing in its skin color. The phenotype of ‘CG’ was initially observed in Talagante (Metropolitan Region, Chile) during grape harvest in 2005, within a cv. ‘RG’ field. ‘Chimenti Globe’ has reddish grape clusters and was selected by its producer for propagation via cuttings (www.chgchile.cl).

## MATERIALS AND METHODS

### Plant material and experimental design

Plant material used in this study corresponds to six plants of commercial table grape cv. ‘Red Globe’ (RG) and six plants of cv. ‘Chimenti Globe’ (CG), positioned in Camino Loreto, Parcela 8-9, Talagante, (Región Metropolitana, Santiago of Chile; 33°38′26′′S; 70°51′52′′ W) (FIGURE 1A). Sampling was performed during seasons 2013 and 2014. Veraison stage was set as the period at which clusters were 30–50% colored and 5° Brix (5% w/w soluble solids) (FIGURE 1B), and the late ripening stage when berry sugar content reached 22-23°Brix (FIGURE 1C). One cluster from each plant (experimental unit) was randomly sampled in both cultivars, in the two stages and seasons. Berries were peeled, and berry skins were frozen in liquid nitrogen and stored at –80°C until required.

**FIGURE 1:**
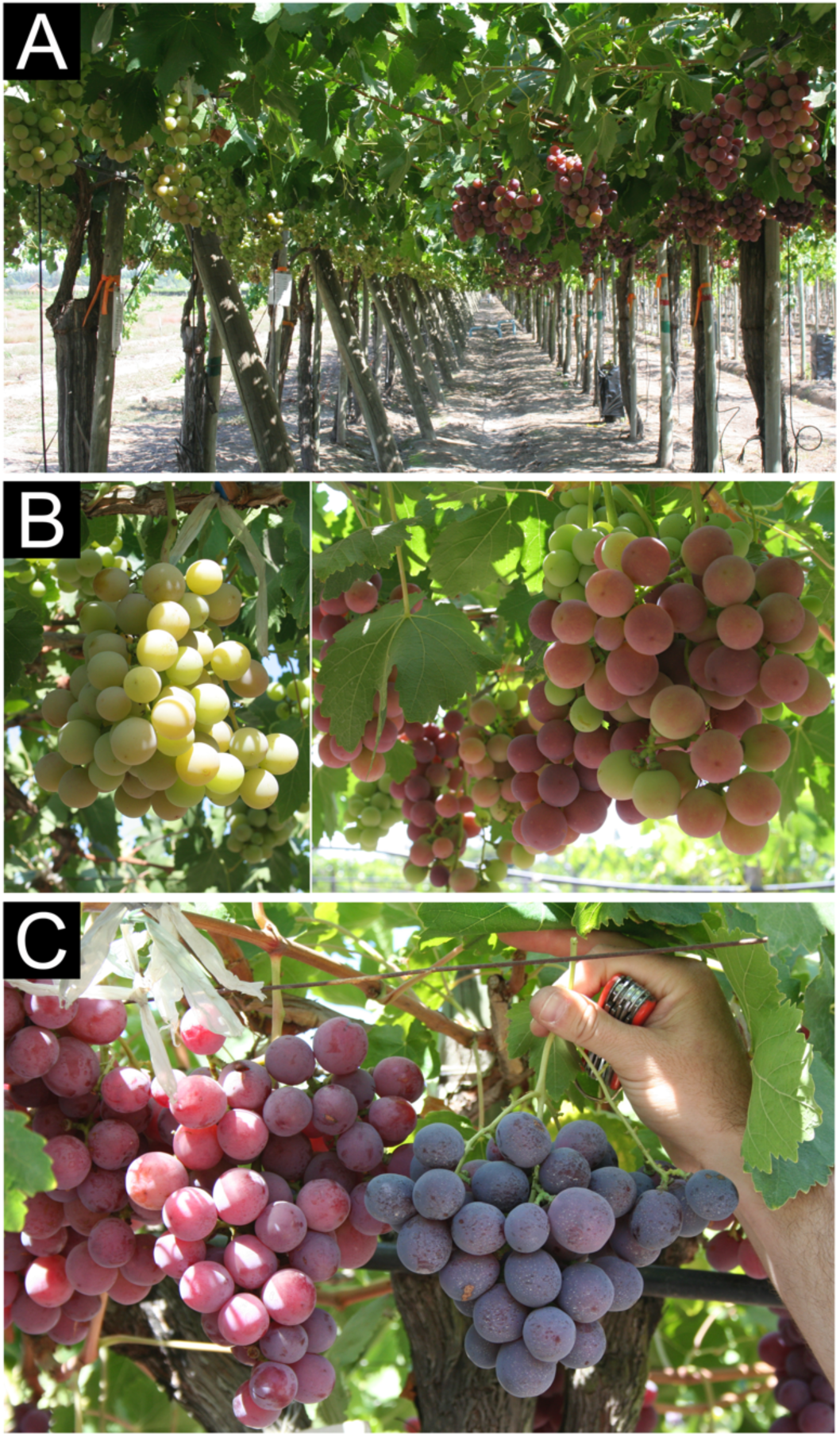
Pigmentation differences at field of the color somatic variants under study. **(A-B)** Cultivar (cv.) ‘Chimenti Globe’ (CG) and cv. ‘Red Globe’ (RG) at the onset of ripening/veraison (EL-35) and **(C)** at a late ripening stage (EL-38). Clusters of CG plants are shown at the left side of each photograph in all cases. Plants from both cultivars used in this study are situated immediately adjacent to each other in the same field.

### Microscopy

Oblique hand sections of ripening berries were cut with a scalpel blade and floated on a glycerol 50% droplet and mounted on slices. Samples were examined under spectral confocal optical microscope *Eclipse C2si*, Nikon Instruments Inc. (Minato-ku, Tokyo, Japan).

### Metabolite extractions and quantification analyses

For metabolite analyses, berry skins from both seasons were used and processed independently (24 total samples per cultivar), except for the case of anthocyanin analysis, for which five out of six samples were considered for each cultivar (20 samples per cultivar). Primary metabolites such as Trehalose-6-Phosphate (T6P) together with glycolytic and tricarboxylic acid (TCA) cycle intermediates were extracted from 25 mg of frozen powder with chloroform/methanol solution, and measured by high-performance anion-exchange liquid chromatography coupled to tandem mass spectrometry (LC-MS/MS) as described by (John E. Lunn et al., 2006). Amino acids were extracted and analyzed by HPLC as described by (Martínez-Lüscher et al., 2014). For sugars and organic acids, an aliquot of 150-200 mg of skin fine powder was extracted sequentially with ethanol 80% and 50%, dried in a SpeedVac and re-dissolved in 1 ml of sterile ultrapure water. Hexose content (glucose and fructose) was measured enzymatically with an automated micro-plate reader (Elx800UV, Biotek Instruments Inc., Winooski, VT, USA) according to (L. Gomez, Bancel, Rubio, & Vercambre, 2007). Tartaric acid content was measured by using the colorimetric method based on ammonium vanadate reactions (Pereira et al., 2006). Malic acid content was determined using an enzyme-coupled spectrophotometric method that measures the change in absorbance at 340 nm from the reduction of NAD^+^ to NADH (Pereira et al., 2006).

Anthocyanins were extracted from 300 mg freeze-dried ground powder from skin of cv. ‘RG’ and ‘CG’ using 1 ml methanol containing 0.1% HCl (v/v). Extracts were filtered across 0.45 μm polypropylene syringe filter (Pall Gelman Corp., Ann Harbor, MI, USA) for high performance liquid chromatography (HPLC) analysis. Each individual anthocyanin was analyzed as described in (Dai et al., 2013). For anthocyanin quantification was integrated peak area at 520 nm and using Malvidin 3-glucoside as standard (Extrasynthèse, Lyon, France).

### RNA extraction

Total RNA were isolated from berry skins according to the procedure of (Reid, Olsson, Schlosser, Peng, & Lund, 2006), using a CTAB-spermidine extraction buffer. This material was used for transcriptome sequencing by RNA-seq technology and validation of data by quantitative real-time RT-PCR (qRT-PCR). For RNA sequencing, berry skins from 2013 season were used and pooled in 3 samples per cultivar (cv. ‘CG’ and ‘RG’) and stage (veraison and ripening), reaching a total number of 12 samples to be deep sequenced. Each pool was generated by mixing 2 individual RNA extractions (1 µg each one) and then mixing them to obtain a concentration of 2 µg per pool (TABLE S1). Total RNA was sent to Macrogen (Macrogen Inc. Seoul, Korea) after ethanol precipitation (0.1 volume of 3 M Sodium Acetate pH 7-8 and 2 volumes of 100% ethanol).

### RNA Sequencing and Read Mapping

Ten micrograms of total RNA were fragmented, converted to cDNA, and amplified by PCR to Illumina ® TruSeqTM RNA Sample Preparation Kit (Illumina, Inc., USA). Pair-end 100bp sequence reads were generated using the Illumina Genome Analyzer II (Illumina) and Illumina HiSeq 2000 (Illumina) at Macrogen Inc. according to the manufacturer’s recommendations. Trimmomatic (Bolger, Lohse, & Usadel, 2014) was used to trim and clip reads prior to mapping, removing the adapter sequences as well as the low-quality sequences from the ends of the reads. All the distinct clean reads were aligned to the Grape Genome Database hosted at CRIBI V2 (http://genomes.cribi.unipd.it/grape/) (Vitulo et al., 2014). Uniquely mapped reads were counted by HITSAT2 software (Kim, Langmead, & Salzberg, 2015) and featureCounts in the Rsubread package (Liao, Smyth, & Shi, 2013).

### Analysis of differentially expressed genes (DEGs)

DESeq2 was used for determining differentially expressed genes (DEGs) using a false discovery rate (*FDR*) threshold of 0.05 and an absolute value log_2_ ratio ≥ 1 (Love, Huber, & Anders, 2014). MultiExperiment Viewer (MeV) was used for gene clustering analysis that was performed by the k-means method with Euclidean distance. MeV also was used for graphical representation of DEGs in a heatmap using Fold Change ≥ 1 (Howe, Sinha, Schlauch, & Quackenbush, 2011). Additionally, we performed a new DESeq2 analysis with the same parameters described previously but removing the filter from the Fold Change; the DEGs obtained with this method were annotated using Mercator web tool and then loaded into MapMan software (Lohse et al., 2014; Usadel et al., 2009).

### Validation of RNA-seq data by quantitative real-time RT-PCR (qRT-PCR)

Two micrograms (ug) of total RNA were treated with TURBO DNA-freeTM DNase (Ambion®) and subsequently reverse transcribed with random hexamer primers and SuperScript II RT (InvitrogenTM Co., Carlsbad, CA, USA) as in (Dauelsberg et al., 2011). Relative transcript quantification of differentially expressed genes (DEGs) was performed by real-time RT-PCR (qRT-PCR) using the BRILLIANT II SYBR® GREEN QPCR Master Mix and the Mx3000P qPCR system (Stratagene, Agilent Technologies Inc., Santa Clara, CA, USA) according to the manufacturer’s instructions. Expression levels of all evaluated genes were calculated from six biological replicates, relative to *Vvi60SRP* housekeeping control gene. We used the 2^-ΔΔCt^ method for the statistical analysis (Schefe, Lehmann, Buschmann, Unger, & Funke-Kaiser, 2006). Primers are listed in Supplementary (TABLE S2).

### Statistical data analyses

Data were analyzed with multivariate analysis methods using the R statistics environment (R Core Team, 2010). In order to evaluate alterations in metabolite levels during stages of grapevine development, principal component analysis (PCA) was performed on mean-centered and scaled data using the ade4 package in R (Dray & Dufour, 2007). Differences between developmental stages and seasons were analyzed by a two-way ANOVA followed with a Tukey multiple comparison post-hoc test at *P* < 0.05, for example, in the case of PCA analysis, component 1. For differences between cultivars were analyzed using unpaired T-test, for example in the case of PCA analysis, component 2 and qRT-PCR analysis. GraphPad Prism version 6.00 for Windows (GraphPad Software, La Jolla California USA) was used for graphics representation and analysis. For RNA-seq analysis, Trimmomatic (Bolger et al., 2014), HISAT2 (Kim et al., 2015) and Rsubread package (Liao et al., 2013) were used with default parameters. DEGs analysis were performed with DEseq2 using *FDR* with a threshold of 0.05 and absolute value log_2_ ratio ≥ 1 (Love et al., 2014). (MeV) was used for cluster analysis by the k-means method with Pearson’s correlation distance and graphical representation of DEGs with a heatmap using FC ≥ 1 (Howe et al., 2011).

## RESULTS

### Characterization of cv. ‘Red Globe’ and its color somatic variant cv. ‘Chimenti Globe’

#### Microscopy study

Grape berry skin is composed of several cell layers: the epidermal cells comprising only a single layer (L1-derived) and the large underlying subepidermal cells (L2-derived) that also compromise the flesh. We observed in cv. ‘Chimenti Globe’ that anthocyanins only accumulated in the outermost single layer of the Epidermis (Ep) and not in the subEpidermis (subEp) (FIGURE 2A-B-C). A clear accumulation of anthocyanins was observed within the vacuoles of the epidermal cells, in Anthocyanin Vacuolar Inclusions (AVIs) (FIGURE 2C). This observation was similar to the previously characterized cv. ‘Malian’ (a bud sport of cv. ‘Cabernet Sauvignon’; Walker et al., 2006) and cv. ‘Pinot Gris’ (a periclinal chimera of cv. ‘Pinot Noir’; Vezzulli et al., 2012). In contrast, cv. ‘Red Globe’ showed colored cells in the Ep layer (FIGURE 2E), with diffuse edge, and some specific AVIs inside some subEp cells that form part of the skin (FIGURE 2D).

**FIGURE 2:**
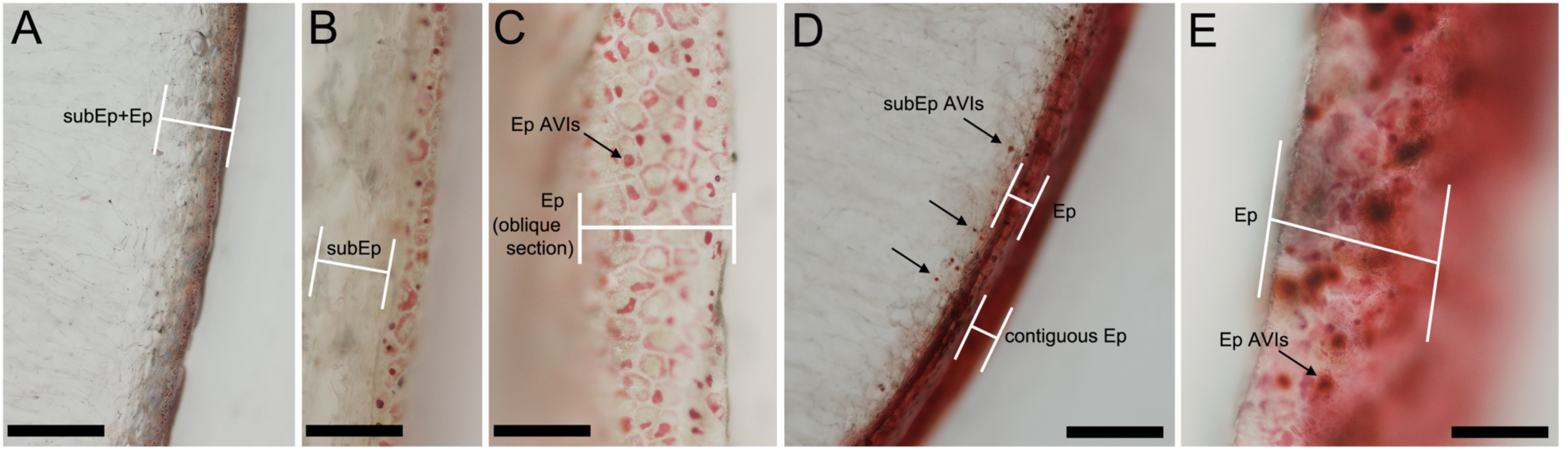
Differential accumulation of anthocyanins in epidermal and subepidermal cell layers of CG and RG. Light microscopy sections showing cells from the epidermis (Ep; L1 ontology) and subepidermis (subEp; L2 ontology) in cv. ‘Chimenti Globe’ **(A-C)** and cv. ‘Red Globe’ **(D-E)**. Anthocyanin vacuolar inclusions (AVIs) are depicted with arrows. Oblique angle hand sections permitted observe contiguous cells from the epidermis as in Walker et al. (2006). In both cultivars the mesocarp/flesh (L2 ontology) is devoid of anthocyanins. Bar in A and D is 500 μm; B, C and E is 100 μm

#### Primary metabolite profiling identifies several compounds affected in the skin of ‘CG’

The composition of the main primary metabolites was assessed by different methods depending of each metabolite (described in Methodology). Quantifications were analyzed with Principal Component Analysis (PCA) to obtain a comprehensive view of main metabolites showing differences between both somatic variants in veraison and ripening. The first two principal components (PC_1_ and PC_2_) explained about 64% of the total variance and allowed to discriminate developmental stages between CG and RG (FIGURE 3A). PC_1_ (43,75%) was inferred to capture predominantly variation according to developmental stage and also to the effect of season but the latter exclusively for the ripening stage samples (no variation was observed for season at the veraison stage). Metabolites contributing to these differences were related to phenylpropanoid metabolism such as shikimate, UDP-glucose and phenylalanine but in addition the molecular regulator trehalose-6-phosphate (T6P) and TCA cycle intermediates such as citrate, isocitrate and several amino acids also contributed to differentiate veraison from ripening (FIGURE 3B and FIGURE S1). PC_2_ variation (20,26%) was associated to cultivar type, but this discrimination was much more evident for the ripening stage samples. These results are explained in changes observed in metabolites related with biosynthesis of sucrose: glyceraldehyde 3-phosphate, fructose 6-phosphate, glucose 6-phosphate, fructose 1,6-biophosphate, glucose 1,6-biophosphate, glycerol 3-phosphate, fructose, glucose and phosphoenolpyruvate; and also, with anthocyanin compounds (FIGURE 3B and FIGURE S2).

**FIGURE 3:**
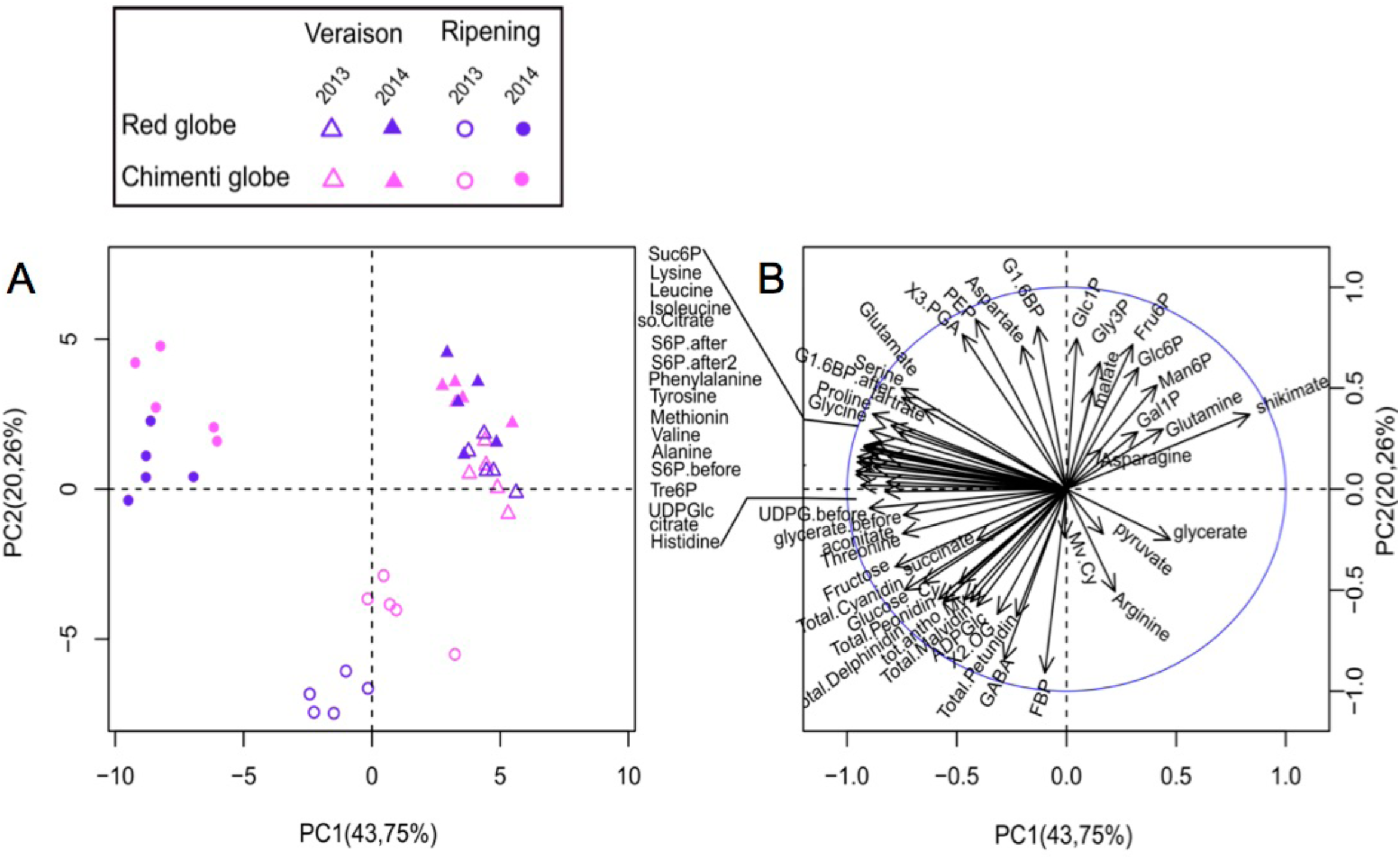
Differential metabolomes in berries of color somatic variants influenced by developmental stage and vintage. (A) Principal component analysis (PCA) of primary metabolites present in berry skins of RG and CG, discriminating by cultivar depending on the developmental stage. The variance explained by season (2013/2014) is implied at both stages but is more evident at the ripening stage. (B) Loading plots of metabolites analyzed for the first two components (PC1 and PC2). Main metabolites that explained PC1 correspond to those close to the ordinate axis such as tre6P, UDPGlc, methionine, phenylalanine and shikimate; and for PC2 correspond to those close to the abscissa axis such as phosphorylate intermediates and anthocyanins.

#### Anthocyanin profiles in berry

We analyzed anthocyanin levels present in berry skins of cv. ‘RG’ and its somatic variant ‘CG’ at the same developmental stages and seasons described previously. During the veraison stage, ‘CG’ showed trace amounts of some anthocyanin types (Table S3). At ripening, we observed the most significant differences in total anthocyanin abundances between cv. ‘CG’ and ‘RG’, with RG containing more than seven times higher amount (FIGURE 4A).

**FIGURE 4:**
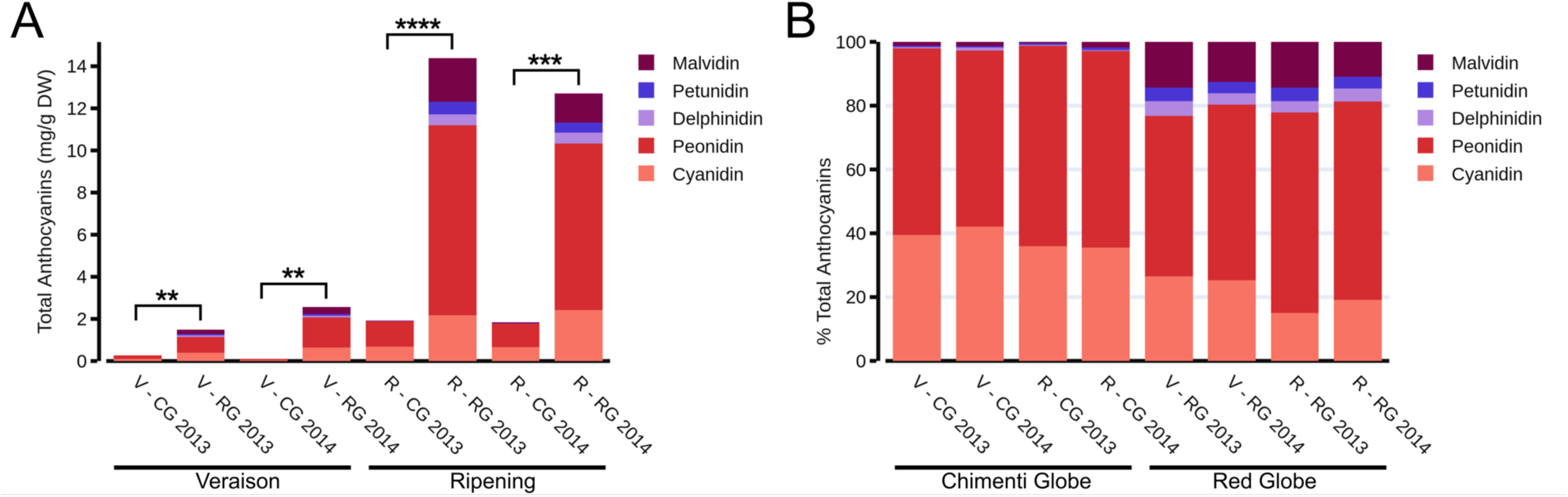
The decreased anthocyanin content in cv. ‘Chimenti Globe’ is accentuated by the insufficiency in accumulating tri-hydroxylated derivatives. (A) Content of total anthocyanins (mg g^-1^ DW, all derivatives) in both somatic variants at veraison and ripening stages in 2013 and 2014 seasons. Asterisks indicate statistical significance as the result of an unpaired T-test. **, *P* < 0.0081; ***, *P* = 0.0001; ****, *P* < 0.0001.). (B) Relative abundance of each anthocyanin type for each sample. Di-hydroxylated derivatives found were cyanidin 3-glucoside, cyanidin 3-(*p*-coumaryl)-glucoside, peonidin-3-glucoside and peonidin-3-(*p*-coumaryl)-glucoside. Tri-hydroxylated derivatives found were delphinidin 3-glucoside, delphinidin 3-(acetyl)-glucoside, petunidin 3-glucoside and malvidin-3-glucoside. Values in (A) and (B) represent the mean of five biological replicates. Anthocyanin derivative content for each replicate can be found in Supplementary Table S3). DW: Dry Weight.

Detailed analysis of anthocyanin derivatives revealed that ‘CG’ mainly contains di-hydroxylated types, namely peonidin (in the form of peonidin 3-glucoside and peonidin 3-(*p*-coumaryl)-glucoside) and cyanidin (in the form of cyanidin 3-glucoside and cyanidin 3-(*p*-coumaryl)-glucoside), which confer more reddish tones (FIGURE 4B) compared to tri-hydroxylated forms. Tri-hydroxylated anthocyanins (the most commonly and abundantly found in black-skinned *V. vinifera* cultivars and that confer purplish and bluish colors), were present in very small amounts or even non-detectable in cv. ‘CG’, while they were found increasing towards ripening in cv. ‘RG’. The proportion of total tri-hydroxylated derivatives with respect to that of di-hydroxylated forms was maintained relatively constant between veraison and ripening (between 20-35% of the total amount of anthocyanins), being Malvidin the most abundant of all three (TABLE S3).

### Genome-wide characterization of cv. ‘Chimenti Globe’ and ‘Red Globe’ transcriptomes

#### RNA-Seq data mapping to the grapevine genome and category enrichment analysis

We originally hypothesized that the genes responsible for changes in berry skin color in cv. ‘CG’ should be differentially altered in expression when compared to cv. ‘RG’. We performed mRNA-Seq profiling using twelve RNA samples corresponding to three independent pools of biological replicates for ‘CG’ and ‘RG’ for each developmental stage in the 2013 season (TABLE S1). From a total range of ∼4.8 – 8.1 million sequencing raw read pairs, from each pool sample, only ∼4.5 – 7.6 million read pairs corresponded to both kept reads that passed the quality analysis, representing a 93% of the total read pairs (TABLE S4). Approximately ∼3.3 –5.1 million reads aligned uniquely to the annotated transcriptome, while ∼0.5 – 1.5 million reads aligned 0 times and ∼0.7 – 1.2 million reads aligned multiple times (72%, 11-20% and 15-17% of the total reads after quality analysis, respectively) (TABLE S5). Only the reads that aligned uniquely and where mapped to the genome were further considered in the analysis (TABLE S5, S6).

To explore the differences between both color somatic variants, all clean reads were aligned to the grape genome database V2 (Vitulo et al., 2014) and then a comparative transcriptomic analysis was performed to screen for differentially expressed genes (DEGs) using DEseq2 package (Love et al., 2014). The first inspection of the complete RNA-seq dataset showed that the major differences between both somatic variants are observed at ripening (FIGURE 5). The PC1, accounting for approximately 50% of the total variance, was able to discriminate ‘CV’ and ‘RG’ only at ripening while the PC2 accounted for differences mostly attributable to ‘RG’ at ripening compared to the three other samples (with the exception one replicate in CG ripening).

**FIGURE 5:**
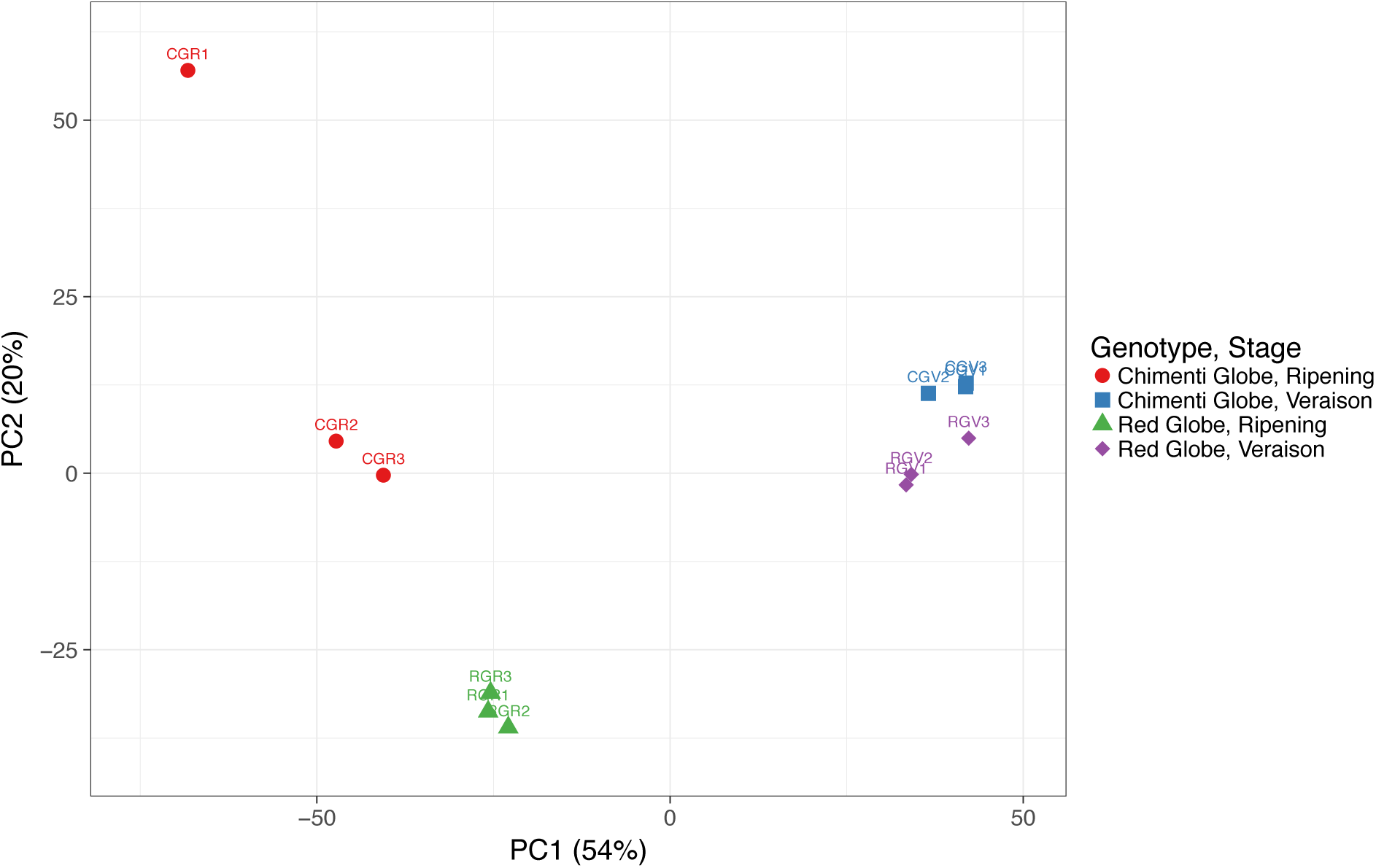
PCA of transcriptomics data. X and Y axis show principal component 1 and principal component 2 that explain 54% and 20% of the total variance, respectively. Original FPKM values are ln(x + 1)-transformed (NAs removed). No scaling was applied to individual genes and SVD with imputation was used to calculate principal components. N = 12 data points.

A total of 438 genes showed differential expression (log2FC>1 and p adjusted value<0.05) in at least one stage. We performed a Gene Ontology (GO) enrichment analysis to enlighten functional categories represented in the 438 DEGs (Supplemental Dataset 1), by using the Mercator web tool and then loaded into MapMan software version 3.5.1R2 (Lohse et al., 2014; Usadel et al., 2009). Three major types of secondary metabolic pathways were differentially expressed between cultivars at veraison and ripening: terpenoids and the phenylpropanoid-related lignin/lignans and flavonoids (FIGURE S3). Additional categories found enriched were those related to enzymes families (e.g. cytochrome P450, UDP-glycosyltransferases, glutathione-S-transferases, glucosidases, o-methyltransferases, peroxidases and beta 1,3 glucan hydrolases, FIGURE S4), and those related to biotic stresses (FIGURE S5) including genes involved in the synthesis of stress-related hormones, pathogenesis related (PR) proteins and transcription factors, among others.

#### The cultivar ‘Chimenti Globe’ exhibits an altered expression of the entire flavonoid pathway compared to cv. ‘Red Globe’

From the 438 DEGs, a total of 109 genes were differentially expressed at both veraison and ripening stages. These were grouped in 6 clusters according to a hierarchical clustering of their expression profiles (FIGURE 6). From the 109 DEGs, many were related to flavonoid metabolism. Cluster A consisted in only one gene: a *glutathione-S-transferase* collinear with GST4 (VIT_204s0079g00710) that is involved in anthocyanin transport in grapes. Cluster B (23 genes) and cluster C (31 genes), where genes are down-regulated in cv. ‘CG’ at both developmental stages, contained many flavonoid/anthocyanin-related genes such as GST13 (VIT_207s0104g01800), GST4 (VIT_204s0079g00690), *flavonoid 3,5-hydroxylases* (F3’5’-H, VIT_206s0009g02805 – 2810 – 2830 – 2840 – 2846 – 2860 – 2873 – 2880 – 2970 – 2920 – 3010), *anthoMATEs (*AM1, VIT_216s0050g00900 and AM2, VIT_216s0050g00910*), flavanone 3-hydroxylase* (F3-H, VIT_204s0023g03370)*, anthocyanin membrane protein 1 (*ANM1, VIT_208s0007g03560*), anthocyanin acyltransferase (*3AT, VIT_203s0017g00870) *chalcone synthase* (CHS, VIT_205s0136g00260), anthocyanin-o-methyltranferases (AOMT1, VIT_201s0010g03510 and AOMT2, VIT_201s0010g03490) and *UDP-glucose:flavonoid 3-O-glucosyltransferase* (UFGT, VIT_216s0039g02230). These clusters also included the genes *vacuolar invertase 1* (GIN1, VIT_216s0022g00670) involved in sugar transport, *cellulose synthase (*CSLA02, VIT_206s0061g01230) and an uncharacterized *cytochrome b5* (CYTB5, VIT_218s0001g09400) (FIGURE 6). Another interesting gene repressed in cv. ‘CG’ in both developmental stages encodes a *homeobox-leucine zipper* (HD-Zip) transcription factor (VIT_216s0100g00670) homologue to *anthocyaninless 2,* a homeobox gene involved in anthocyanin distribution and root development in Arabidopsis (Kubo, Peeters, Aarts, Pereira, & Koornneef, 1999) (FIGURE 6). Cluster E (34 genes) encompassed genes that were induced at both developmental stages in cv. ‘CG’, some of which may be related to phenylpropanoid metabolism, such as *cytochrome p450* (CYP71E1, VIT_201s0137g00540), *ABC transporter family protein* (VIT_214s0060g00720) and a *catechol-O-methyltransferases* (VIT_215s0048g02480-2490). The transcription factor *MYB24* (VIT_214s0066g01090) and a *sesquiterpene synthase* (TPS10, VIT_218s0001g04780) were also listed in this group. Cluster D (11 genes) and Cluster F (9 genes) combined genes with contrasting expressions at veraison and ripening. Genes falling in these categories included three (VIT_210s0042g00840, VIT_210s0042g00880 and VIT_210s0042g00920) and *ABC transporter G member 15* (VIT_209s0002g03560) that were induced at ripening but repressed in veraison (FIGURE 6).

**FIGURE 6:**
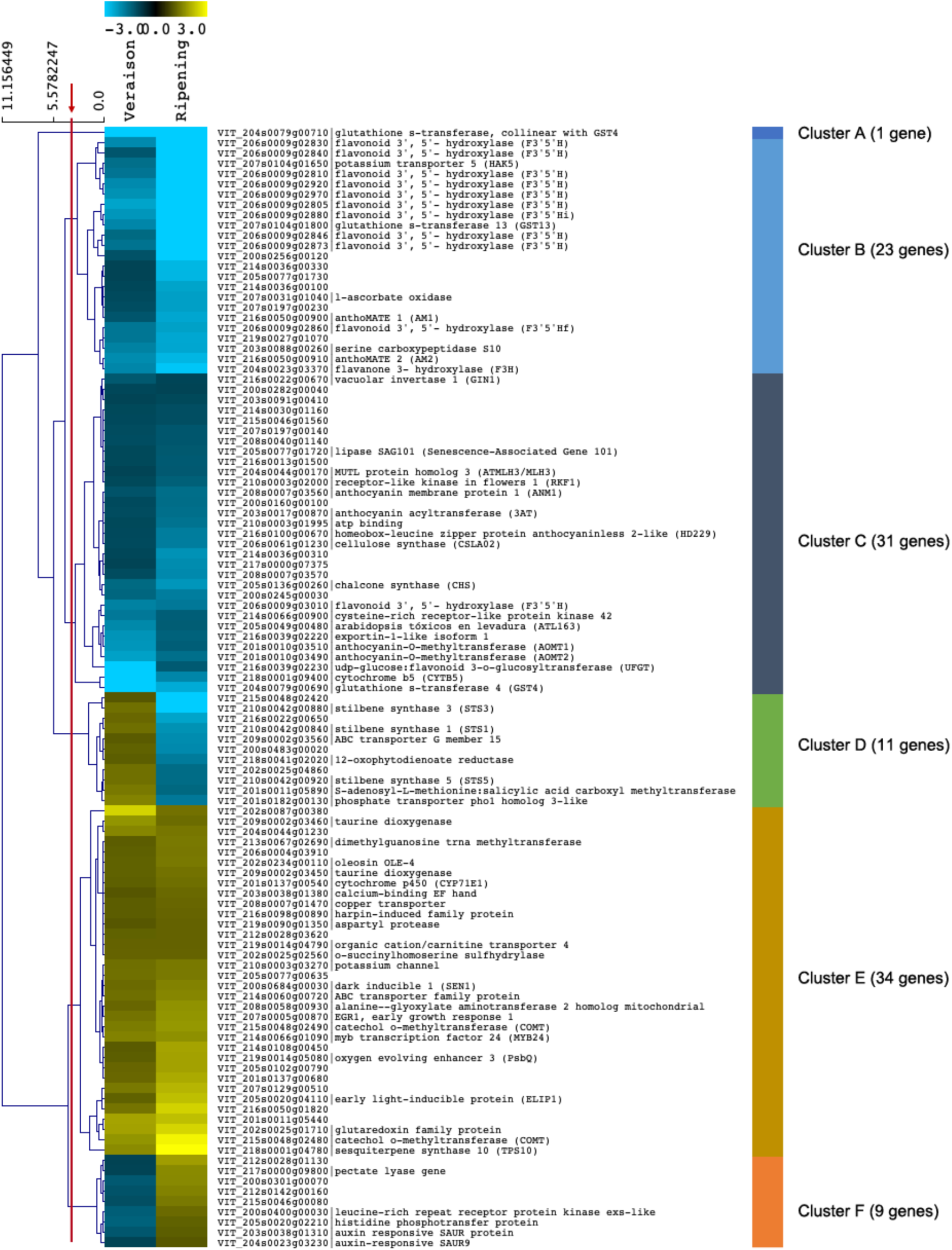
Differentially expressed genes (DEGs) associated to somatic color variation at both veraison and ripening as surveyed by RNA-seq. Heatmap of 109 DEGs in CG compared to RG, where differential expression is defined by logFC>=1 of the CG/RG ratio and FDR<=0.05. Hierarchical clustering shows the presence of six major clusters outlined on the right panel (these were defined by cutting the dendrogram at the site shown by the red line). 12X.v1 (CRIBIv2 annotation) gene IDs were included for those genes with either an annotation or those previously characterized in grape with relevant functions to our study.

We performed a less stringent analysis excluding the Fold Change filtering step to search for additional pathways altered by color somatic variation. Within transcription factors, we found that *MYBA1* (VIT_202s0033g00410), *MYBA8* (VIT_202s0033g00380), *MYB30B* (VIT_214s0108g00830), *MYB141* (VIT_214s0108g01080) and *MYB147* (VIT_217s0000g09080) were repressed at veraison in cv. ‘CG’, while the stilbene regulator *MYB15* (VIT_205s0049g01020), *MYB24* (VIT_214s0066g01090) and *MYB162* (VIT_219s0090g00590) were induced at this stage. The truncated and non-functional *MYBA3* gene (VIT_202s0033g00450, collinear with *MYBA1* and *MYBA2*), and *MYB24* were also induced at ripening. In contrast to what was observed at veraison, *MYB30B* was induced during ripening in cv. ‘CG’. Furthermore, the flavonoid regulator *MYBPA1* (VIT_215s0046g00170) and *MYB145* (VIT_219s0014g03820) were repressed during the ripening stage in cv. ‘CG’. Additional TF genes were differentially expressed in ripening cv. ‘CG’ berries, such as bHLH 145-like gene (VIT_218s0001g07210) and an uncharacterized WD repeat-containing protein C16727 isoform 1 (VIT_208s0007g00730), both being induced (Supplemental Dataset 2).

#### Genes related to ‘Primary metabolism’

Inspection of the less stringent list of DEGs allowed to identify several genes related to primary metabolism between cv. ‘CG’ and cv. ‘RG’. At veraison, we identified genes related to sucrose and trehalose 6-phosphate biosynthesis and also to sugar transport, pathways that have been related in *Arabidopsis thaliana* to a homeostatic mechanism of maintaining sucrose levels within a range that is appropriate for each cell type and developmental stage of the plant (Tre6P:sucrose ratio; Yadav et al., 2014). The bidirectional sugar transporters *VviSWEET1* (VIT_218s0001g15330), *VviSWEET10* (VIT_217s0000g00830) and *VviSWEET15* (VIT_201s0146g00260), three hexose transporters (VIT_205s0020g03140, VIT_211s0149g00050 and VIT_214s0006g02720) and two genes encoding a fructose-bisphosphate aldolase (VIT_203s0038g00670 and VIT_204s0023g03010) were induced in cv. ‘CG’ in the veraison stage. At ripening, we observed genes related to sucrose and trehalose biosynthesis and sugar transportation such as *VviSWEET11* (VIT_207s0104g01340, induced in cv. ‘CG’), two genes encoding a sucrose-phosphate synthase (VIT_218s0075g00330 and VIT_218s0089g00490, induced and repressed at ripening, respectively), a sucrose transporter and sucrose synthase 1 (VIT_201s0026g01960 and VIT_207s0005g00750) both being repressed. A fructose-1,6-bisphosphatase (VIT_208s0007g01570) and two genes putatively encoding a fructose-bisphosphate aldolase (VIT_204s0023g03010 and VIT_203s0038g00670) were induced. Finally, four genes putatively encoding a trehalose-phosphate synthase (VIT_206s0009g01650, VIT_217s0000g08010, VIT_203s0063g01510 and VIT_212s0028g01670) were induced in cv. ‘CG’ during ripening (Supplemental Dataset 3).

#### Genes from other secondary metabolic pathways

Other genes found in the less-strict list were related to other metabolic pathways such as isoprenoid biosynthesis: two isoprene synthase genes up-regulated in cv. ‘CG’ at veraison (VIT_212s0134g00020, VIT_212s0134g00030) and ripening (VIT_212s0134g00030); and the carotenoid pathway: a probable carotenoid cleavage dioxygenase 4 gene down-regulated in cv. ‘CG’ during veraison (VIT_202s0087g00930). Amongst DEGs in cv. ‘CG’, the terpenoid biosynthesis pathway genes TPS35 (VIT_212s0134g00030) and TPS10 (VIT_218s0001g04780) were up-regulated both at veraison and ripening.

#### Validation of the RNA-seq gene expression profiles

Twelve DEGs were selected for qRT-PCR validation (FIGURE 7). From clusters of down-regulated genes (clusters A, B and C) nine genes were selected: nine isoforms of *F3’5’-H*, *CYTB5*, *UFGT*, *AOMT1-2*, *AM1-2*, *GST13*, *F3H* and *CHS*. Seven of these genes (*F3’5’-H*, *CYTB5*, *UFGT, AOMT1-2, GST13, F3-H* and *CHS*) showed significant differences between cv. ‘CG’ and cv. ‘RG’ at veraison and ripening, fully supporting the RNA-seq results, while *AM1-2* only showed differences at the ripening stage (FIGURE 7).

**FIGURE 7:**
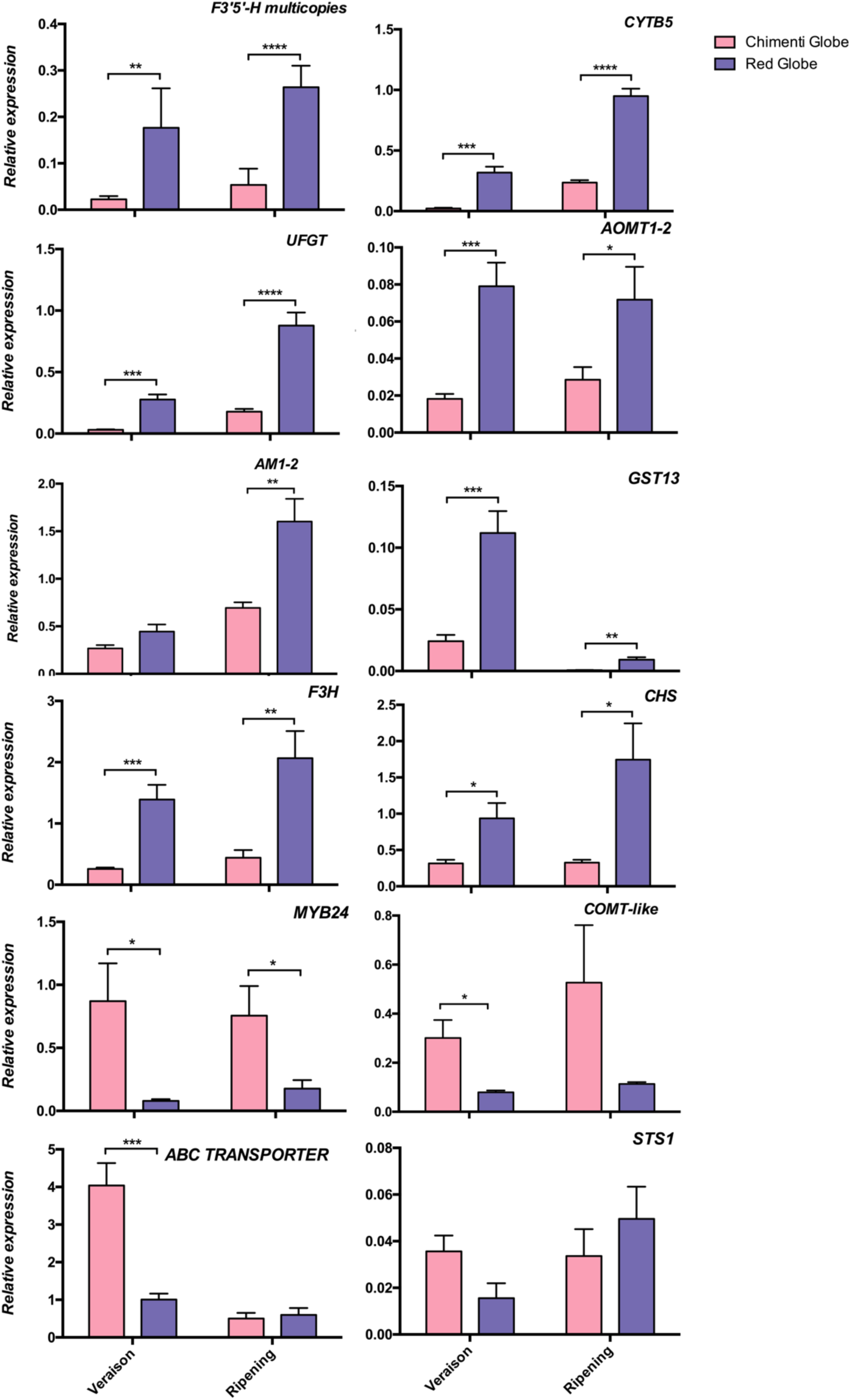
Real-time quantitative PCR (qRT-PCR) validation of selected DEGs in berry skins of CG and RG. Samples were collected at veraison and ripening stages in 2013. Genes were selected based in their role or position in the repression, induction or combined repression/induction clusters. Gene expression in berries is shown relative to *SRP60* (VIT_205s0077g02060) housekeeping gene expression. Values represent the mean of six biological replicates, and bars represent the standard error of mean. These were derived from one qPCR run with duplicate PCR reactions on each of the biological replicates. Gene expressions were normalized together for both developmental stages. Asterisks indicate statistical significance as the result of unpaired T-test between both somatic variants, independently taken at each developmental stage (*, *P* < 0.0465; **, *P* < 0.0053; ***, *P* < 0.0009; ****, *P* < 0.0001). FW, Fresh Weight.

To gain more insight in gene expression of the nine isoforms of *F3’5’-H* we were able to design specific primers for six of them. In general, we observed significant differences in each isoform (i.e. induction in cv. ‘RG’ compared to cv. ‘CG’) at both stages (FIGURE S6B). We confirmed this observation by inspecting the specifically-assigned reads in the RNA-seq analysis (TABLE S7). From the Cluster of induced genes (Cluster E), three were further selected, validating *MYB24*’s significant induction in both veraison and ripening stages, while an *ABC transporter family protein* (VIT_214s0060g00720) and two *COMT-like* genes (designed primers recognized both VIT_215s0048g02480 and VIT_215s0048g02490) were induced in cv. ‘CG’ only at veraison (FIGURE 6). The *STS1* gene, that codified a Stilbene synthase was selected from clusters D, exhibiting a higher expression at ripening and a lower one at veraison in cv. ‘CG’ compared to cv. ‘RG’, but without significant differences.

We observed that within the list of genes belonging to the repression cluster on the heatmap generated (FIGURE 6) there were 11 copies of the gene *F3’5’-H* not induced in cv. ‘CG’: VIT_206s0009g02805, VIT_206s0009g02810, VIT_206s0009g02840, VIT_206s0009g02860, VIT_206s0009g02873, VIT_206s0009g02880, VIT_206s0009g02920, VIT_206s0009g02970 and VIT_206s0009g03010. These genes were located closely in the same region of chromosome 6 (FIGURE S6A). This corresponded to a segment of the chromosome that span from approximately 15,660 to 16,060 MB.

### Exploration of the MYBA color locus identifies MYBA1 as the only gene responsible for the skin color differences

In order to try to elucidate the genetic origin of the ‘Chimenti Globe’ phenotype, we checked the presence of MYBA1 white and red alleles in cv. ‘CG’ and cv. ‘RG’ using the same set of primers used in (Shimazaki et al., 2011), while also studied *MYBA1* expression levels by qPCR. We observed the presence of white and red alleles of *MYBA1* in all cv. ‘CG’ and cv. ‘RG’ samples tested (FIGURE 8), demonstrating that both cultivars are heterozygous for this gene, at least in the L2 cell layer. These results were validated with molecular marker analysis (TABLE S8), which in addition revealed a homozygous null allele configuration of MYBA2, in both L1 and L2 cell layers, in the two somatic variants. We further confirmed the null allele of MYBA2 in >99% samples (both cultivars) by inspecting all the reads matching the G/T positions related to MYBA2 mutation (i.e. variant calling analysis; TABLE S9). As MYBA2 is non-functional in both cultivars, only the differences in *MYBA1* expression could account for the skin pigmentation differences of ‘Chimenti Globe’. We tested *MYBA1* expression by qRT-PCR, observing that it was down-regulated at veraison (RNA-seq log_2_FC −0,947) and ripening (confirmed only by qRT-PCR) in cv. ‘CG’ (FIGURE 8).

**FIGURE 8:**
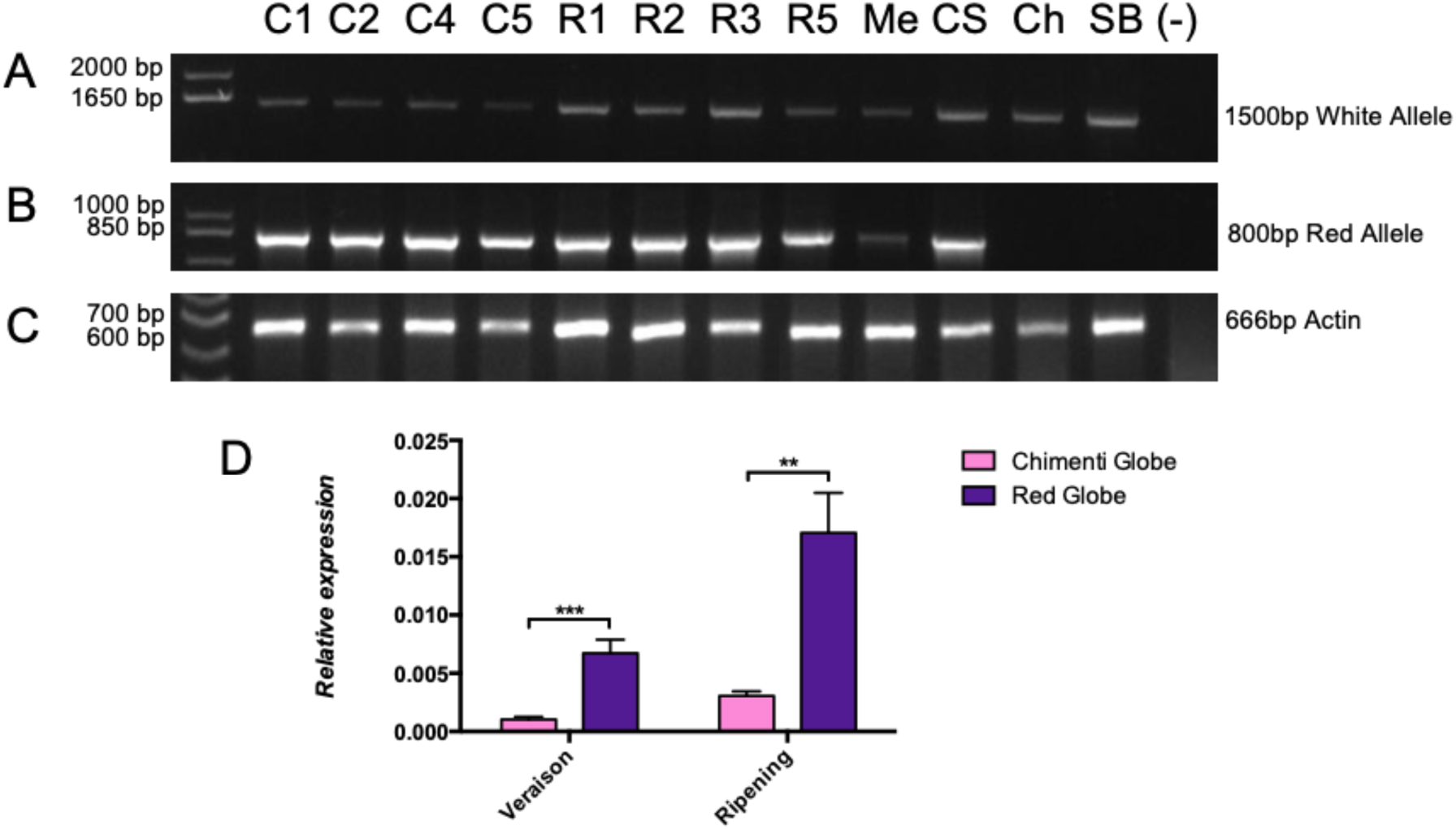
Heterozygous *VviMYBA1* gene allelic configuration in both somatic variants and reduced *MYBA1* expression in cv. ‘Chimenti Globe’ compared to cv. ‘Red Globe’. (A-B) PCR identification of the (A) non-functional white allele and (B) functional red allele in cv. ‘Chimenti Globe’ (C1,2,3), cv. ‘Red Globe’ (R1,2,3), and other cultivars used as controls (cv. ‘Merlot’ -Me, cv. ‘Cabernet Sauvignon’ -CS, cv. ‘Chardonnay’ -Ch and cv. ‘Sauvignon Blanc’ -SB); (C) *ACTIN* gene amplification was used as an internal PCR control check for gDNA integrity. Numbers on the left side of each electrophoresis gel photograph indicate the band size according to the DNA marker ladder, while numbers on the right show the exact size of the white and red alleles according to (Shimazaki, Fujita, Kobayashi, & Suzuki, 2011). (D) qRT-PCR analysis of *VviMYBA1* in berry skins. Relative gene expression calculation as the mean of six biological replicates, with error bars representing the standard error of the mean. Asterisks indicate the result of unpaired T-test. **, *P* = 0.0023; ***, *P* = 0.0007.

## DISCUSSION

### Anthocyanin accumulation in the berry skin of cv. ‘Chimenti Globe’ is diminished in correlation to the repression of both *MYBA1* and the flavonoid/anthocyanin pathways

Grape somatic variants have been well described in the literature and specifically those affecting skin pigmentation have become a source of new commercial genotypes (Azuma, 2018; Pelsy, 2010; Torregrosa et al., 2011; Vezzulli et al., 2012; Walker et al., 2006; Carbonell-Bejerano et al., 2017). In this work, we studied cv. ‘Chimenti Globe’ (‘CG’) that was originated as a somatic variation of cv. ‘Red Globe’ (‘RG’). Microscopic analysis showed in ripe berries that cv. ‘CG’ had a particular pattern of anthocyanin accumulation, exclusive to the epidermis. In contrast, ‘RG’ also presented pigmentation in the sub-epidermis. This pattern was similarly described in cv. ‘Malian’ (Walker et al., 2006) and cv. ‘Pinot Gris’ (Vezzulli et al., 2012). The microscopy analysis allowed to suggest initially that cv. ‘CG’ was a periclinal chimera with only the L1 cell layer capable of producing anthocyanins. However, the molecular marker analysis showed that ‘CG’ was heterozygous for the functional *MYBA1* allele in the L2. Under this configuration, MYBA1 is still active and transcriptionally regulates the accumulation of anthocyanins in vacuoles (Ford, Boss, & Hoj, 1998). In the case of pink-skinned cultivars of oriental origin, Shimazaki et al. (2011) reported a 33bp insertion in the second intron of *VviMYBA1* as a consequence of low expression levels of *VviMYBA1*. Thus, it would be necessary to sequence *MYBA1*’s genomic region in both L1 and L2 derived tissues and see if there is any change that may account for *MYBA1* lower expression in ‘CG’.

The lower *VviMYBA1* expression, together with the particular pattern of anthocyanin accumulation in cv. ‘CG’ observed by microscopy i.e. absence of anthocyanin vacuolar inclusions in subepidermal cells, allows to suggest anthocyanin transport as a possible process being largely affected and responsible for diminished anthocyanin accumulation. In plant cells, anthocyanins are synthesized in the cytoplasmatic face of the endoplasmic reticulum (Winkel-Shirley, 2001; Winkel, 2004) and then are transported to the vacuole for storage. Three different mechanisms have been described for flavonoid transport: membrane transporters, vesicle trafficking and glutathione S-transferase (GST)-mediated transport (Zhao, 2015). In grapevine, orthologs of TT12 have been identified, an Arabidopsis gene encoding a MATE protein required for anthocyanin and proanthocyanidin accumulation in the vacuole (Marinova et al., 2007). In addition, AM1 and AM3 transport acylated anthocyanins in the presence of Mg ATP (C. Gomez et al., 2009); and *VviABCC1* transports malvidin 3-O-glucoside into vacuoles (Francisco et al., 2013). The glutathione-S-transferases *VviGST1*, *VviGST3* and *VviGST4*, expressed in grape fruits, have the ability to bind different flavonoid ligands (Pérez-Díaz, Madrid-Espinoza, Salinas-Cornejo, González-Villanueva, & Ruiz-Lara, 2016). From the 109 differentially expressed genes from our study, we observed several genes related to anthocyanin vacuolar transportation being repressed in cv. ‘CG’: *GST* and two *anthoMATE* (AM1 and AM2) genes. Interestingly, *VviGST4 and VviAM2* are specifically highly expressed at berry maturity (Sun et al., 2016). Other genes down-regulated in cv. ‘CG’ were *ANM1*, encoding a putative anthocyanin membrane protein 1, and *3AT* related with anthocyanin acylation, a process known to improve anthocyanin intensity and color stabilization (Yonekura-Sakakibara, Nakayama, Yamazaki, & Saito, 2008). All these genes are directly regulated by MYBA1 and in less intensity by its homologues MYBA6 and MYBA7 (Matus et al., 2017)

### Metabolic changes during veraison and ripening stages in cv. ‘Red Globe’ and cv. ‘Chimenti Globe’

Berry ripening is an active process that involves regulation of global gene expression (Fasoli et al., 2012) and changes in the accumulation of primary and secondary metabolites (Boss, Davies, & Robinson, 1996; Conde et al., 2007). Primary metabolites and in particular sugars are indeed elicitors of anthocyanin synthesis, reason why we decided to analyze them in this study. Metabolic comparison between the two somatic variants showed that the developmental stage was the primary discriminant parameter, despite the use of different cultivars. This observation is concordant with previous published studies (D.-L. Guo et al., 2016; Massonnet et al., 2017; D. C. J. Wong, Lopez Gutierrez, Dimopoulos, Gambetta, & Castellarin, 2016; Wu et al., 2014) in which, despite the use of different cultivars, the importance of developmental stages appeared to overrule cultivar origin. T6P seems a main influential metabolite (as seen in PC1, FIGURE 3A). T6P is a sugar phosphate considered as a metabolic signaling molecule in plants that regulates developmental growth (John Edward Lunn, Delorge, Figueroa, Van Dijck, & Stitt, 2014) and was demonstrated to be involved in sugar signaling in *Arabidopsis thaliana* (John E. Lunn et al., 2006). In *Vitis vinifera* there is no clear evidence of control of T6P during berry development, although several transcriptomic studies have shown that orthologue genes encoding T6P synthase, T6P synthase/phosphatase and SnRK1 (Sucrose-non-fermentative Related Kinase 1, i.e. another sugar signaling pathway component) are tightly regulated during berry development (Deluc et al., 2007; Gambetta, Matthews, Shaghasi, McElrone, & Castellarin, 2010). Although no significant differences were observed at the level of quantification of T6P, the analysis obtained from the RNA-seq analysis data (DEG analysis without Fold Change filter) showed the down-regulation of four genes encoding trehalose - phosphate synthase in cv. ‘CG’ during ripening).

Due to the fact that T6P modulates sugar signaling, we searched for DEGs between cv. ‘RG’ and cv. ‘CG’ related with sugar, fructose, glucose and sucrose biosynthesis in our RNA-seq analysis. We identified three genes at veraison, *VviSWEET1* (VIT_218s0001g15330), *VviSWEET10* (VIT_217s0000g00830) and *VviSWEET15* (VIT_201s0146g00260) and in ripening *VviSWEET11* (VIT_207s0104g01340), as induced in cv. ‘CG’. These constitute a protein family of sugar uniporters involved in sugar export (Chen et al., 2010; Chong et al., 2014; W.-J. Guo et al., 2014). Indeed, it has been described that several SWEET proteins in *V. vinifera* increase from veraison onwards in cv. ‘Syrah’ and cv. ‘Muscatel Ottonel’ (Chong et al., 2014). Despite these genes are less expressed in green berries, *VviSWEET1* is the mainly expressed in young and adult leaves; *VviSWEET10* and *VViSWEET11* in flowers; *VviSWEET10*, *VviSWEET11* and *VviSWEET15* in berries after veraison (Chong et al., 2014).

We found other genes related directly with sucrose biosynthesis induced in cv. ‘CG’: two fructose-bisphosphate aldolases (VIT_203s0038g00670 and VIT_204s0023g03010) were induced during veraison and ripening stage. This enzyme catalyzes the conversion of glyceraldehyde 3-phosphate (PGAL) to fructose 1,6-bisphosphate (F1,6-BP). Concordantly, we also found that these metabolites were decreased in cv. ‘CG’ during ripening stage in 2013, and in fact was one of the metabolites that explained the differences observed in cultivars (PC2, FIGURE 3A). A similar pattern was observed with fructose-1,6-bisphosphatase (VIT_208s0007g01570), induced in cv. ‘CG’. This enzyme catalyzes the conversion of F1,6-BP to fructose 6-phosphate (F6-P), which was found decreased in cv. ‘CG’ at ripening. The sucrose-phosphate synthases encoded by VIT_218s0075g00330 and VIT_218s0089g00490 catalyze the conversion of uridine diphosphate-glucose (UDP-glucose) to glucose-fructose-phosphate (GF-P), leading to the last step of sucrose biosynthesis (Ruan, 2014).

Taking all these observations together, it is possible to suggest an alteration in cv. ‘CG’ occurring at the level of sugar export and in sucrose biosynthesis. This process could be related to the anthocyanin biosynthesis pathway in cv. ‘CG’ as anthocyanin accumulation in plants can be induced by sugar (Das, Shin, Choi, & Park, 2012). Specifically in grapevines, fructose, glucose and sucrose induce anthocyanin accumulation (Dai et al., 2013; Larronde et al., 1998; Zheng et al., 2009).

Among primary metabolites occurring in the intersection with secondary metabolism, we found shikimate as significant. This is an intermediary of the shimikate pathway that mediates the carbon flow from carbohydrate metabolism to phenylpropanoid and aromatic compound biosynthesis (Maeda & Dudareva, 2012; Zhang et al., 2012). This strongly suggest the importance of this metabolite in species such as grapevine, in which the berry consumes phenylalanine for the biosynthesis of flavonoid compounds such as anthocyanins.

Anthocyanins explained major differences in the PC analysis of metabolites. We observed that only traces of malvidin, petunidin and delphinidin were detected in cv. ‘CG’, indicating a block in the tri-hydroxylated branch of the anthocyanin biosynthesis pathway, an observation latter confirmed by the RNA-seq analysis. Accordingly, an increase in F3’5’-H activity has been previously reported to explain higher concentration of tri-hydroxylated anthocyanins in Cabernet Sauvignon and Shiraz, pinpointing F3’5’-Hs as key molecular players driving the phenylpropanoid metabolic flux towards bluish anthocyanin production (Degu et al., 2014).

### Transcriptomic changes during veraison and ripening in cv. ‘CG’ and cv. ‘RG’

In our study, besides metabolic studies, the skin transcriptomes of both somatic variants were analyzed using Illumina Hiseq2000. We report that several genes related to flavonoid pathway, grouped in three main clusters, were differentially expressed in cv. ‘CG’ compared to cv. ‘RG’. Among these, we identified eleven *F3’5’-H* isoforms repressed in cv. ‘CG’, consistent with the depletion of delphinidin-type anthocyanins. The adjacent location of each one of these copies in a single cluster within chromosome 6 (FIGURE S6A; Castellarin et al., 2006; Falginella et al., 2010) allows to suggest a common regulation for all of them, presumably by MYBA1 and other TFs. *CYTB5* was down-regulated in cv. ‘CG’ at both stages. *CYBT5* has been previously involved in F3’5’-H activity modulation in Petunia flowers, catalyzing the transfer of electrons to their prosthetic heme group (de Vetten et al., 1999). In 2006, (Jochen Bogs, Ebadi, McDavid, & Robinson, 2006) demonstrated that a putative *CYBT5* from grapevine cv. ‘Shiraz’ modulated both F3’5’-H and F3’-H activities even though the exact mechanism remains to be deciphered. This is of interest because this suggests a possible link regarding the drastic down-regulation of eleven gene copies of F3’5’-H.

Three *Stilbene synthase* genes (*STS1, STS3* and *STS5*), involved in resveratrol biosynthesis (Dubrovina & Kiselev, 2017; Kiselev, Aleynova, Grigorchuk, & Dubrovina, 2016) were induced in cv. ‘CG’ at veraison and down-regulated during ripening stage. MYB15 is involved in the transcriptional regulation of stilbene synthases in grapevine (Holl et al., 2013) and was also induced in cv. ‘CG’. This suggests a possible compensation point in flavonoid metabolism: CHS utilizes the same substrates as STS but, is responsible for flavonoid-type compound formation. Chalcone synthase (*CHS*) was indeed down-regulated in cv. ‘CG’, in correlation with less total anthocyanins. Thus, STS could act as an escape valve to reflux the exceeded carbon after the decrease in CHS activity. Taken together, this would suggest a multilevel alteration of the flavonoid pathway, and not only of its anthocyanin tri-hydroxylated branch in cv. ‘CG’.

### The berry color locus shows similar allelic configurations in both somatic variants

White skinned grape cultivars that do not produce anthocyanins contain a *MYBA1* null allele in homozygosity, in which the presence of the retrotransposon *Gret1* in the promoter/5’UTR region interrupts the normal transcription (Kobayashi et al., 2004). Pigmented cultivars possess at least one functional ‘red berry’ allele either without *Gret1* or with the excised retrotransposon (This et al., 2007). We observed the presence of white and red alleles of *MYBA1* in all cv. ‘CG’ and cv. ‘RG’ samples tested, thus, it seems that the allelic configuration of *MYBA1* is not responsible for the phenotype differences. Despite this, the down-regulation of flavonoid and anthocyanin structural genes in cv. ‘CG’ is only possibly related to MYBA1, as MYBA2 appears non-functional in all meristematic cell layers and cultivars. The significant differences of *MYBA1* expression between both cultivars at veraison and ripening should be the responsible of the changes in anthocyanin abundance (including the changes in di/tri-hydroxylated anthocyanin profiles). The expression of *MYBA1* in ‘Chimenti’ is decreased by a different process not identified here.

## AUTHOR CONTRIBUTIONS

EG, PAJ, SD and JTM designed experiments. CS performed material sampling, RNA extraction, for deep sequencing and qRT-PCR, gathered all the results and wrote the manuscript. TM and CM conducted the bioinformatics analyses and data mining. JL and RF carried out the LC-MS/MS metabolic analysis. GH and CR assisted in hexose, anthocyanin and organic acids quantifications. LM assisted in RNA extraction and material sampling collection. RA and DC performed molecular marker analysis. JTM reanalyzed RNA-seq data, produced PCA plots, edited figures and together with FMN, EG, PAJ and SD revised the manuscript.

## ACKNOWLEDGMENTS

We thank to CONICYT scholarship doctorate 2012 N° 21120432, Operational Expenses Scolarship of CONICYT N° 21120432, Millennium Nucleus of Plant Systems and Synthetic Biology NC130030, FONDECYT 1150220 and CHIMENTI S.A. This work was also supported by Grant PGC2018-099449-A-I00 and by the Ramón y Cajal program grant RYC-2017-23645, both awarded to J.T.M. from the Ministerio de Ciencia, Innovación y Universidades (MCIU, Spain), Agencia Estatal de Investigación (AEI, Spain), and Fondo Europeo de Desarrollo Regional (FEDER, European Union).

