## Supplemental Figures and Tables for "Differences in berry primary and secondary metabolisms identified by transcriptomic and metabolic profiling of two table grape color somatic variants"

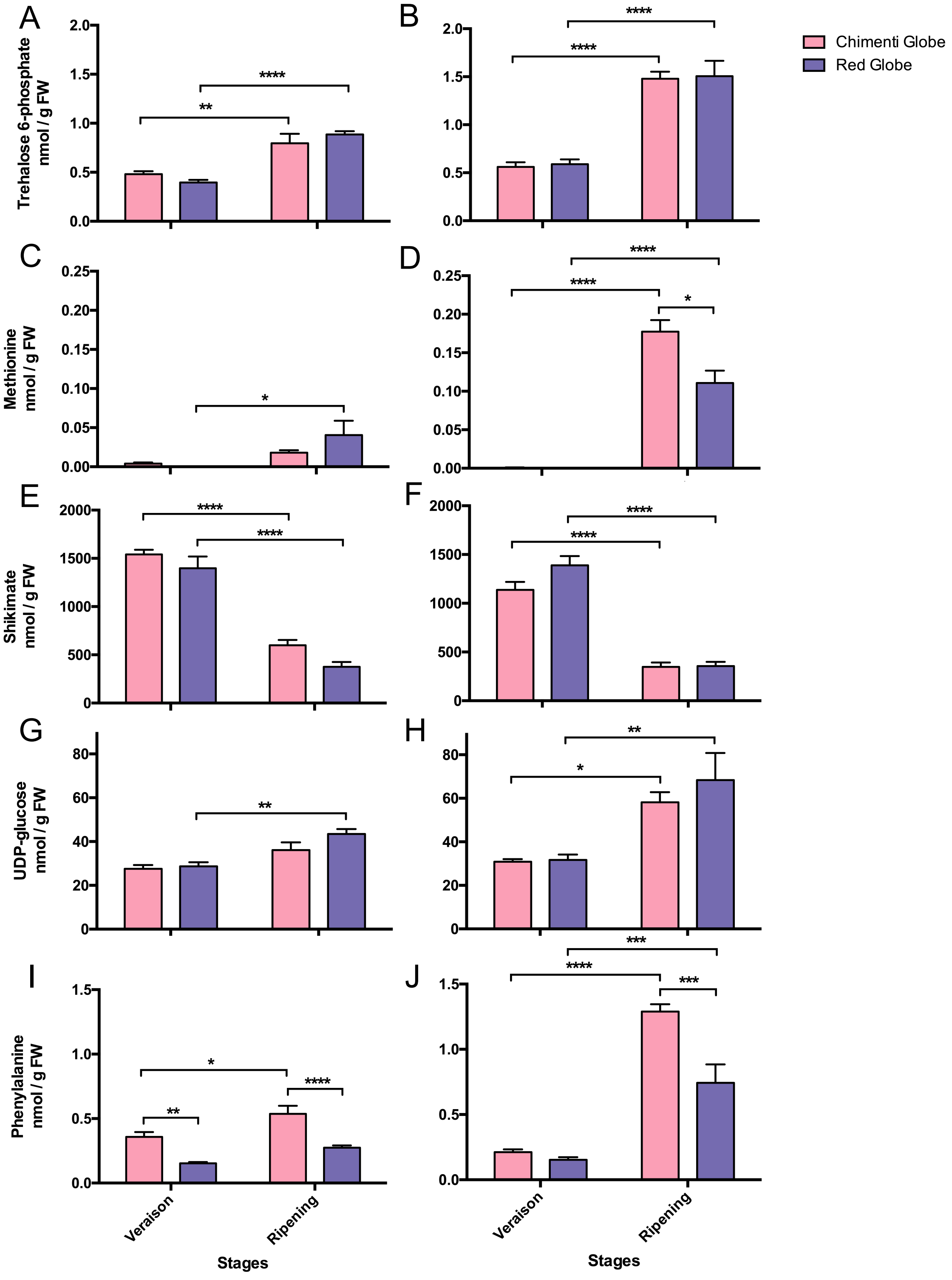
A
B
C
D
E
F
G
H
I
J
Chimenti Globe
Red Globe
Trehalose 6-phosphate nmol / g FW
Methionine nmol / g FW
Shikimate nmol / g FW
UDP-glucose nmol / g FW
Phenylalanine nmol / g FW
Veraison
Ripening
Stages

**FIGURE S1 (previous page): Metabolites found in berry skins of cv. 'Chimenti Globe' and cv. 'Red Globe' whose percentage of contribution are higher in the PC1 of the Principal Component Analysis shown in Figure 2**. Metabolites corresponding to the seasons 2013 and 2014 are shown in **(A, C, E, G, I**) and **(B, D, F, H, J)**, respectively. The abundances reported (nmol $g^{-1}$ FW) correspond to the mean of six biological replicates, with error bars representing standard deviations. Asterisks indicate the result of two-way ANOVA, followed by Tukey's test. ****, $P < 0.0001$; ***, $P < 0.0005$; **, $P < 0.0052$; *, $P < 0,0282$. FW, Fresh Weight.

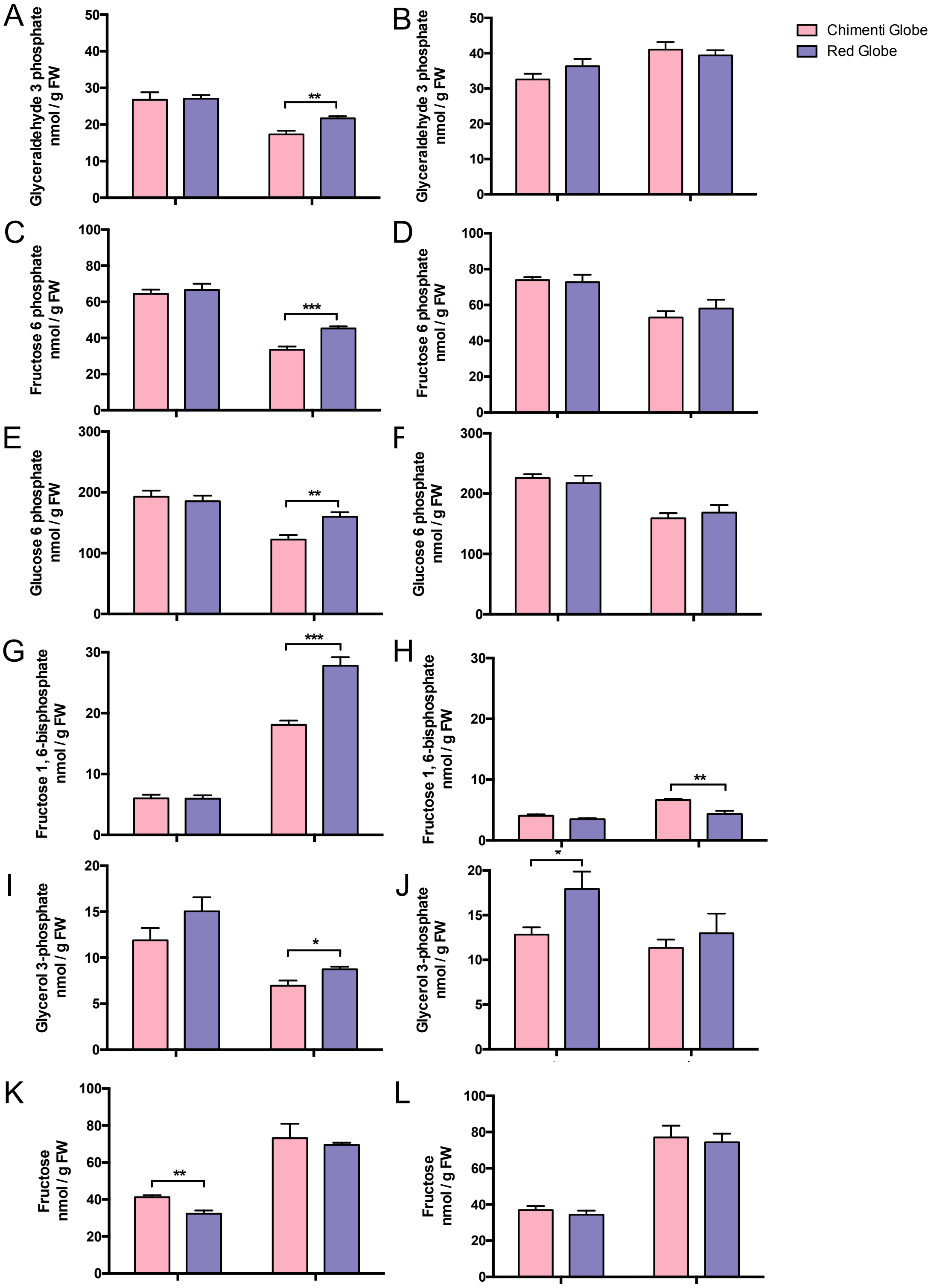
A
B
C
D
E
F
G
H
I
J
K
L
Chimenti Globe
Red Globe
Glyceraldehyde 3 phosphate nmol / g FW
Fructose 6 phosphate nmol / g FW
Glucose 6 phosphate nmol / g FW
Fructose 1, 6-bisphosphate nmol / g FW
Glycerol 3-phosphate nmol / g FW
Fructose nmol / g FW

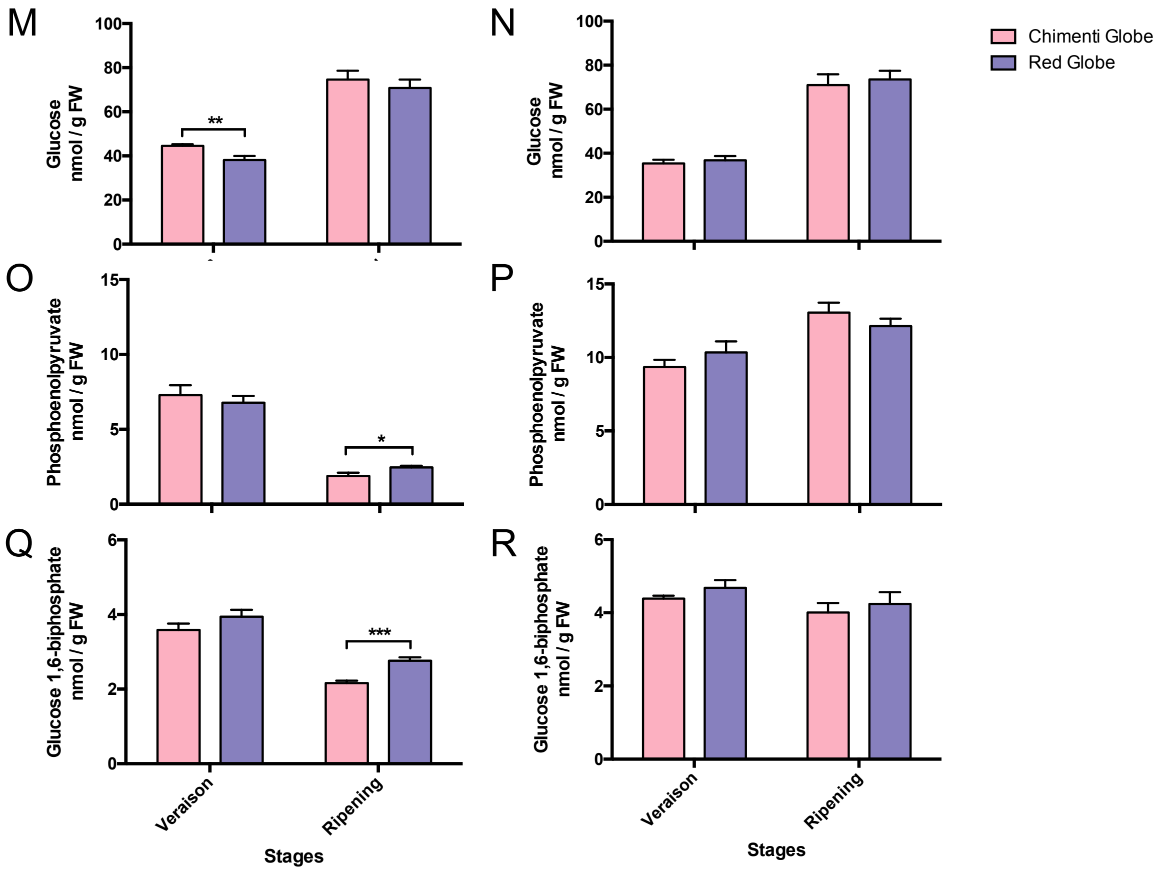


**FIGURE S2: Metabolites found in berry skins of cv. 'Chimenti Globe' and cv. 'Red Globe' whose percentage of contribution are higher in the PC2 of the Principal Component Analysis shown in Figure 2**. Metabolites corresponding to the seasons 2013 and 2014 are shown in **(A, C, E, G, I, K, M, O, Q**) and **(B, D, F, H, J, L, N, P, R)**, respectively. The abundances reported (nmol $g^{-1}$ FW) correspond to the mean of six biological replicates, with error bars representing the standard error from the mean. Asterisks indicate the result of unpaired T-test. *, $P < 0.0443$; **, $P < 0.0085$; ***, $P < 0.0003$.; ****. FW, Fresh Weight.

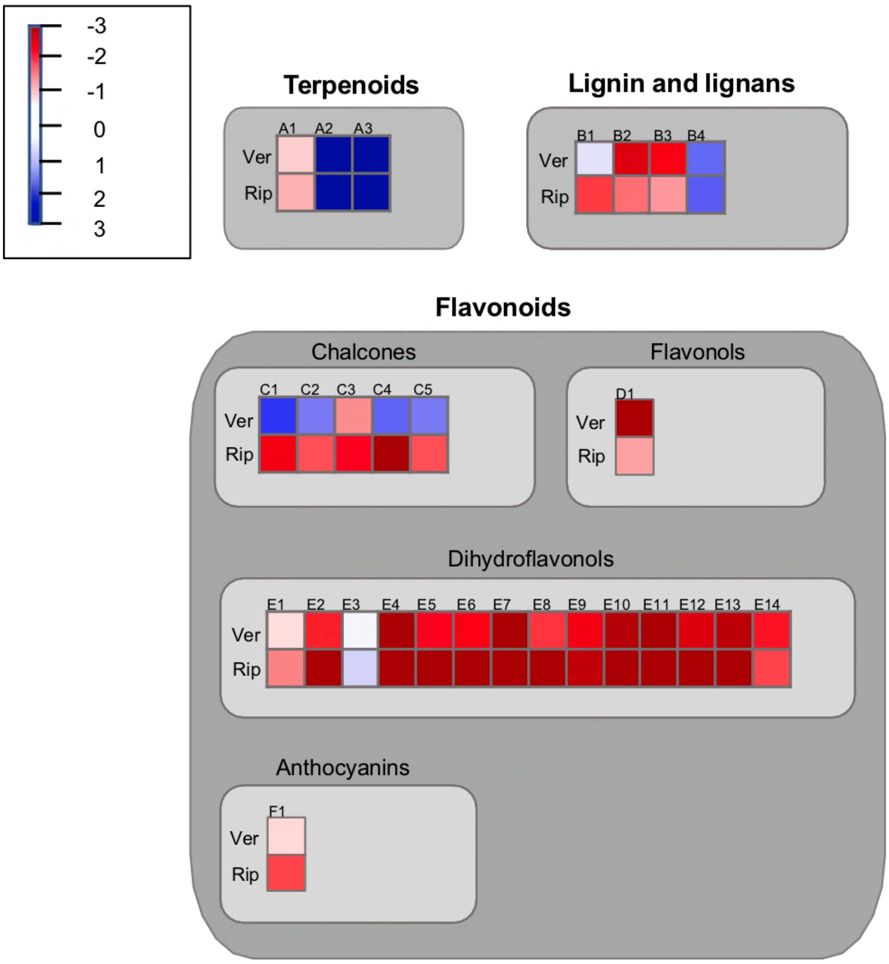


**FIGURE S3: Mapman visualization of gene expression changes related to secondary metabolism observed in cv. 'Chimenti Globe' in comparison to cv. 'Red Globe' found both at veraison and ripening.** A total of 438 genes were found differentially expressed in at least one-time point. Out of these, 462 DEGs were mapped to MapMan category bins as shown in Supplementary Table S8 (some of the data points may be mapped multiple times to different bins). Here, 32 DEGs mapping to secondary metabolism are shown. Each box represents a single gene, named from A to G. The color indicates $log_2$ value of fold change in the CG/RG comparison. Terpenoids belonging to Bincode 16.1.5 (secondary metabolism, isoprenoids, terpenoids): **A1** (VIT_209s0054g01220.1), **A2** (VIT_212s0134g00030.1) and **A3** (VIT_218s0001g04780.1). Ligning and lignans-related genes belonging to Bincode 16.2.1.1 (secondary metabolism phenylpropanoids lignin biosynthesis PAL): **B1** (VIT_213s0019g04460.1); to Bincode 16.2.1.6 (secondary metabolism phenylpropanoids lignin biosynthesis CCoAOMT): **B2** (VIT_201s0010g03490.1) and **B3** (VIT_201s0010g03510.1); and to Bincode 16.2.1.9 (secondary metabolism phenylpropanoids lignin biosynthesis COMT) **B4** (VIT_215s0048g02490.1). Flavonoids/Chalcone-related genes belonging to Bincode 16.8.2 (secondary metabolism flavonoids chalcones): **C1** (VIT_210s0042g00840.1) and **C2** (VIT_210s0042g00920.1); to Bincode 16.8.2.1 (secondary metabolism flavonoids chalcones naringenin-chalcone synthase:) **C3** (VIT_205s0136g00260.1) **C4** (VIT_210s0042g00880.1) and **C5** (VIT_210s0042g00920.1). Flavonoids/Flavonols belonging to Bincode 16.8.4.3 (secondary metabolism flavonoids flavonols flavonol-3-O-rhamnosyltransferase): **D1** (VIT_216s0039g02230.1). Flavonoids/Dihydroflavonols belonging to Bincode 16.8.3.1 (secondary metabolism flavonoids dihydroflavonols dihydroflavonol 4-reductase): **E1** (VIT_218s0001g12800.1); to Bincode 16.8.3.2 (secondary metabolism flavonoids dihydroflavonols flavanone 3-hydroxylase): **E2** (VIT_204s0023g03370.1) and **E3** (VIT_218s0001g14310.1; and to Bincode 16.8.3.3 (secondary metabolism flavonoids dihydroflavonols flavonoid 3"-monooxygenase): **E4** to **E14** (VIT_206s0009g02805.1, VIT_206s0009g02810.1, VIT_206s0009g02830.1, VIT_206s0009g02840.1, VIT_206s0009g02846.1, VIT_206s0009g02860.1, VIT_206s0009g02873.1, VIT_206s0009g02880.1, VIT_206s0009g02920.1, VIT_206s0009g02970.1 and VIT_206s0009g03010.1), respectively. Flavonoids/Anthocyanins belonging to Bincode 16.8.1.1 (secondary metabolism flavonoids anthocyanins leucocyanidin dioxygenase): **F1** (VIT_202s0025g04720.1).

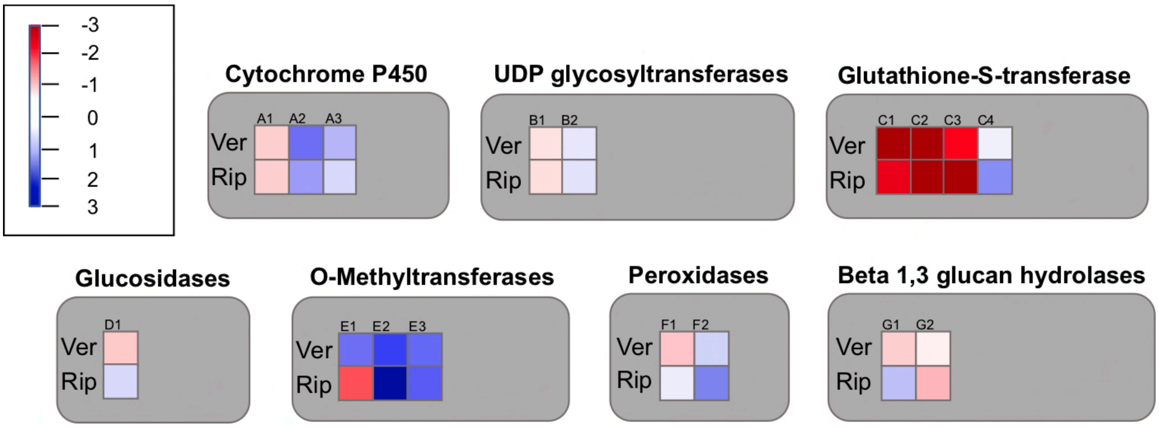


**FIGURE S4: Mapman visualization of gene expression changes related to enzymes families observed in cv. 'Chimenti Globe' in comparison to cv. 'Red Globe' from DEGs found both at veraison and ripening.** Each box represents a single gene named from A to G whose bincode, bin name and more information is in supplemental material (Supplemental Dataset 4). In this visualization were mapped 462 out of 438 DEGs (some of the data points may be mapped multiple times to different bins) and in this pathway were visible 17 DEGs. The color indicates $log_2$ value of fold changes; blue color represents down-regulated transcripts; and red color represents up-regulated transcripts. Cytochrome P450 belonging to Bincode 26.10 (misc cytochrome): **A1** (VIT_200s0705g00010.1), **A2** (VIT_201s0137g00540.1) and **A3** (VIT_202s0025g04850.1). UDP glycosyltransferases belonging to Bincode 26.2 (misc UDP glucosyl and glucoronyl transferases): **B1** (VIT_208s0007g08120.1) and **B2** (VIT_216s0115g00340.1); Glutathione-S-transferase belonging to Bincode 26.9 (misc glutathione S transferases): **C1** (VIT_204s0079g00690.1), **C2** (VIT_204s0079g00710.1), **C3** (VIT_207s0104g01800.1) and **C4** (VIT_218s0001g00690.1); Glucosidases belonging to Bincode 26.3.1 (misc gluco-, galacto- and mannosidases alpha-galactosidase): **D1** (VIT_208s0007g05100.1); O-Methyltransferases belonging to Bincode 26.6 (misc O-methyl transferases): **E1** (VIT_201s0011g05890.1), **E2** (VIT_215s0048g02480.1) and **E3** (VIT_215s0048g02490.1); Peroxidases belonging to Bincode 26.12 (misc peroxidases): F1 (VIT_206s0004g07770.1) and **F2** (VIT_211s0052g00650.1); Beta 1,3 glucan hydrolases belonging to Bincode 26.4.1 (misc beta 1,3 glucan hydrolases): **G1** (VIT_212s0057g00700.1); and to Bincode 26.4.1 (misc beta 1,3 glucan hydrolases glucan endo-1,3-beta-glucosidase): **G2** (VIT_203s0063g02490.1).

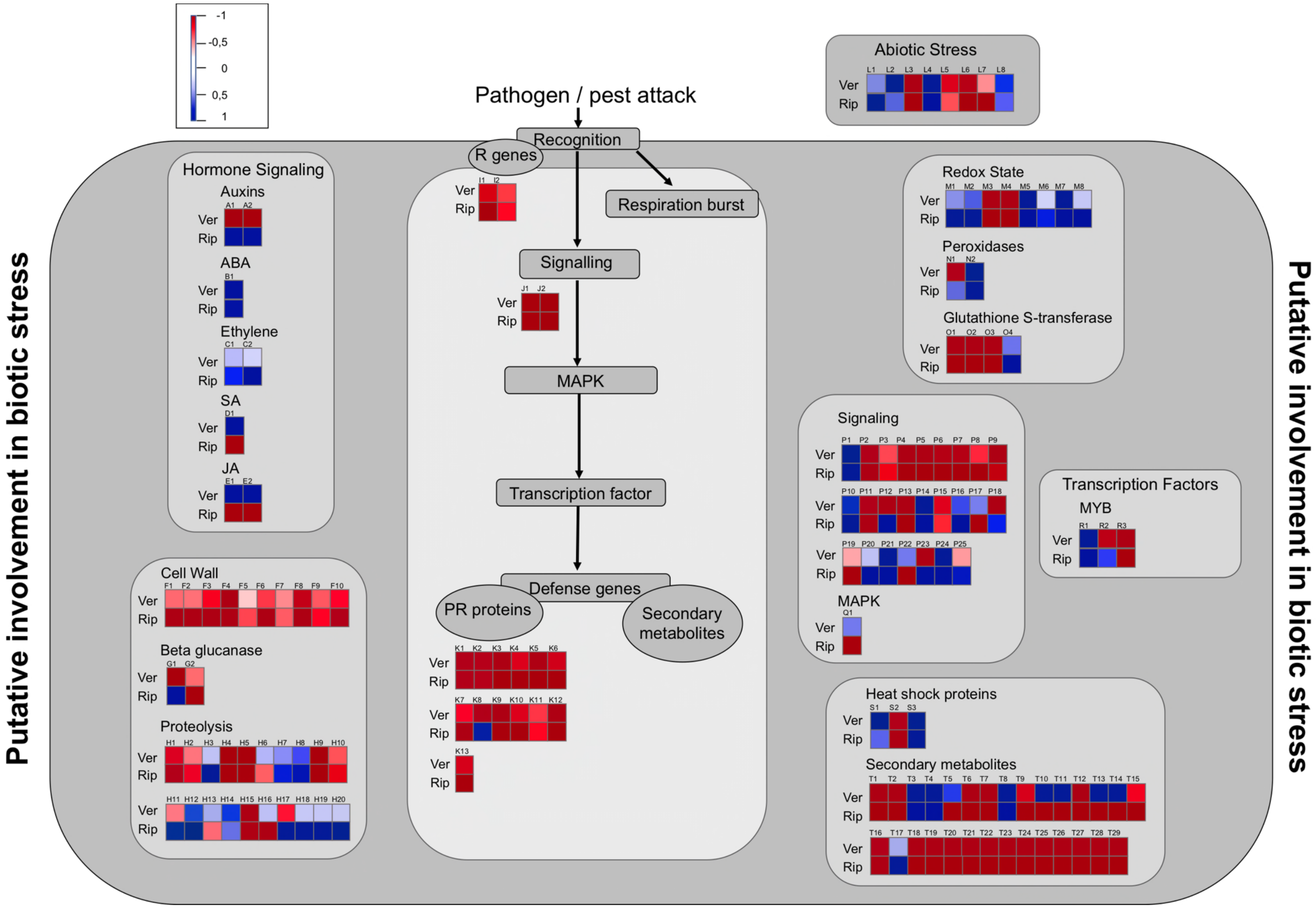
-1
-0,5
0
0,5
1
Pathogen / pest attack
Recognition
R genes
Respiration burst
Signalling
MAPK
Transcription factor
Defense genes
PR proteins
Secondary metabolites
Abiotic Stress
Hormone Signaling
Auxins
ABA
Ethylene
SA
JA
Cell Wall
Beta glucanase
Proteolysis
Redox State
Peroxidases
Glutathione S-transferase
Signaling
MAPK
Transcription Factors
MYB
Heat shock proteins
Secondary metabolites
Ver
Rip
Putative involvement in biotic stress
Putative involvement in biotic stress

**FIGURE S5: Mapman visualization of DEGs related to biotic stress observed in Chimenti Globe in comparison to Red Globe, found both during veraison and ripening stages.** Each box represents a single gene named from A to T whose bincode, binname and more information is in supplemental material (Supplemental Dataset 4). In this visualization were mapped 462 out of 438 DEGs (some of the data points may be mapped multiple times to different bins) and in this pathway were visible 140 DEGs. The color indicates $\log_2$ value of fold changes; blue color represents down-regulated transcripts; and red color represents up-regulated transcripts. Auxins belonging to Bincode 17.2.3 (hormone metabolism auxin induced-regulated-responsive-activated): **A1** (VIT_203s0038g01310.1) and **A2** (VIT_204s0023g03230.1); ABA belonging to Bincode 17.1.3 (hormone metabolism abscisic acid induced-regulated-responsive-activated): **B1** (VIT_217s0000g01080.1); Ethylene belonging to Bincode 17.5.2 (hormone metabolism ethylene signal transduction): **C1** (VIT_205s0049g00510.1) and to Bincode 17.5.3 (hormone metabolism ethylene induced-regulated-responsive-activated): **C2** (VIT_212s0028g02020.1); SA belonging to Bincode 17.8.1 (hormone metabolism salicylic acid synthesis-degradation): **D1** (VIT_201s0011g05890.1); JA belonging to bincode 17.7.1.5 (hormone metabolism jasmonate synthesis-degradation 12-Oxo-PDA-reductase): **E1** (VIT_218s0041g02010.1) and **E2** (VIT_218s0041g02020.1); Cell Wall belonging to 10.1.10 (cell wall precursor synthesis RHM): **F1** (VIT_219s0014g00290.1), to Bincode 10.2 (cell wall cellulose synthesis): **F2** (VIT_200s0580g00010.1), **F3** (VIT_204s0069g00780.1), **F4** (VIT_206s0061g01230.1), to Bincode 10.2.1 (cell wall cellulose synthesis cellulose synthase): **F5** (VIT_205s0020g05050.1), **F6** (VIT_216s0039g02020.1) and **F7** (VIT_218s0122g00120.1), to Bincode 10.6.3 (cell wall degradation pectate lyases and polygalacturonases): **F8** (VIT_205s0051g00590.1) and to Bincode 10.8.1 (cell wall pectin esterases PME): **F9** (VIT_204s0044g01000.1) and **F10** (VIT_204s0044g01010.1); Beta glucanase belonging to Bincode 26.4 (misc beta 1,3 glucan hydrolases): **G1** (VIT_212s0057g00700.1) and to Bincode 26.4.1 (misc beta 1,3 glucan hydrolases glucan endo-1,3-beta-glucosidase): **G2** (VIT_203s0063g02490.1); Proteolysis belonging to Bincode 29.5 (protein degradation): **H1** (VIT_201s0127g00010.1) and **H2** (VIT_206s0061g01180.1), to Bincode 29.5.3 (protein degradation cysteine protease): **H3** (VIT_206s0009g02230.1), to Bincode 29.5.5 (protein degradation serine protease): **H4** (VIT_203s0088g00260.1), **H5** (VIT_208s0040g01140.1) and **H6** (VIT_216s0050g00490.1), to Bincode 29.5.11.3 (protein degradation ubiquitin E2): **H7**

(VIT_214s0108g00140.1), to Bincode 29.5.11.4.2 (protein degradation ubiquitin E3 RING): **H8** (VIT_202s0012g00290.1), **H9** (VIT_205s0049g00480.1), **H10** (VIT_208s0105g00290.1), **H11** (VIT_210s0042g01070.1), **H12** (VIT_211s0065g01210.1), **H13** (VIT_215s0046g01820.1), **H14** (VIT_218s0001g01060.1), **H15** (VIT_218s0001g09530.1), **H16** (VIT_218s0001g11740.1), to Bincode 29.5.11.4.3.2 (protein degradation ubiquitin E3 SCF FBOX): **H17** (VIT_205s0094g01590.1) and to Bincode 29.5.11.20 (protein degradation ubiquitin proteasom): **H18** (VIT_201s0010g02900.1), **H19** (VIT_203s0038g04350.1), **H20** (VIT_210s0042g00070.1); Pathogen/pest attack, R genes belonging to 20.1.2 (stress biotic receptors): **I1** (VIT_218s0089g00100.1) and **I2** (VIT_218s0117g00050.1); Pathogen/pest attack, Signalling belonging to Bincode 20.1.3 (stress biotic signalling): **J1** (VIT_217s0000g07375.1) and **J2** (VIT_217s0000g07400.1); Pathogen/pest attack, PR proteins belonging to Bincode 20.1.7 (stress biotic PR-proteins): **K1** (VIT_200s0160g00100.1), **K2** (VIT_200s0400g00020.1), **K3** (VIT_207s0197g00140.1), **K4** (VIT_207s0197g00170.1), **K5** (VIT_207s0197g00230.1), **K6** (VIT_212s0035g00480.1), **K7** (VIT_213s0067g00910.1), **K8** (VIT_213s0139g00300.1), **K9** (VIT_213s0156g00440.1), **K10** (VIT_218s0089g00100.1), **K11** (VIT_218s0117g00050.1) and **K12** (VIT_219s0027g01070.1), and belonging to Bincode 20.1.7.6.1 (stress biotic PR-proteins proteinase inhibitors trypsin inhibitor): **K13** (VIT_209s0002g00960.1); Abiotic Stress belonging to Bincode 20.2 (stress abiotic): **L1** (VIT_202s0025g04330.1), to Bincode 20.2.1 (stress abiotic heat): **L2** (VIT_206s0004g04470.1), **L3** (VIT_215s0046g01560.1), **L4** (VIT_219s0015g01370.1), to Bincode 20.2.3 (stress.abiotic.drought/salt): **L5** (VIT_201s0026g02340.1), **L6** (VIT_204s0044g01670.1) and **L7** (VIT_218s0001g01270.1) and to Bincode 20.2.99 (stress abiotic unspecified): **L8** (VIT_207s0005g00290.1); Redox State belonging to Bincode 21.1 (redox thioredoxin): **M1** (VIT_204s0023g02700.1) and **M2** (VIT_210s0071g01160.1), to Bincode 21.2 (redox ascorbate and glutathione): **M3** (VIT_218s0001g09400.1), to Bincode 21.2.1 (redox ascorbate and glutathione ascorbate): **M4** (VIT_207s0031g01040.1), to Bincode 21.4 (redox glutaredoxins): **M5** (VIT_202s0025g01710.1) and **M6** (VIT_214s0066g00960.1), belonging to Bincode 21.5 (redox peroxiredoxin): **M7** (VIT_208s0007g03970.1) and to Bincode 21.6 (redox.dismutases and catalases): **M8** (VIT_213s0067g02990.1); Peroxidases belonging to Bincode 26.12 (misc peroxidases): **N1** (VIT_206s0004g07770.1) and **N2** (VIT_211s0052g00650.1); Glutathione-S-transferase belonging to Bincode 26.9 (misc glutathione S transferases): **O1** (VIT_204s0079g00690.1), **O2** (VIT_204s0079g00710.1), **O3** (VIT_207s0104g01800.1) and **O4** (VIT_218s0001g00690.1); Signalling belonging to Bincode 30.1 (signalling. n sugar and nutrient physiology): **P1** (VIT_207s0005g00870.1), to Bincode 30.2.3 (signaling receptor kinases leucine rich repeat III): **P2** (VIT_216s0013g01940.1), to Bincode 30.2.8.1 (signaling receptor kinases leucine rich repeat VIII VIII-1): **P3** (VIT_208s0056g00570.1), to Bincode 30.2.8.2 (signaling receptor kinases leucine rich repeat VIII VIII-2): **P4** (VIT_210s0003g02000.1), **P5** (VIT_212s0055g00590.1), **P6** (VIT_219s0014g00470.1) and **P7** (VIT_219s0014g00620.1), to Bincode 30.2.10 (signaling receptor kinases leucine rich repeat X): **P8** (VIT_200s2085g00010.1) and **P9** (VIT_219s0014g02360.1), to Bincode 30.2.11 (signaling receptor kinases leucine rich repeat XI): **P10** (VIT_206s0004g01750.1), to Bincode 30.2.17 (signaling receptor kinases DUF 26): **P11** (VIT_214s0066g00900.1), to Bincode 30.2.99 (signalling. eceptor kinases misc): **P12** (VIT_205s0020g02210.1) and **P13** (VIT_216s0013g01500.1), to Bincode 30.3 (signaling calcium): **P14** (VIT_203s0038g01380.1), **P15** (VIT_205s0049g01460.1), **P16** (VIT_205s0077g00810.1), **P17** (VIT_207s0031g03030.1), **P18** (VIT_208s0056g00290.1), to Bincode 30.4 (signaling phosphinositides): **P19** (VIT_205s0094g00850.1), to Bincode 30.5 (signaling G-proteins): **P20** (VIT_208s0217g00030.1) and **P21** (VIT_215s0021g01410.1), to Bincode 30.6 (signalling.MAP kinases): **P22** (VIT_200s0567g00010.1), to Bincode 30.10 (signaling phosphorelay): **P23** (VIT_205s0020g02210.1), to Bincode 30.11 (signaling light): **P24** (VIT_205s0020g04110.1), to Bincode 30.11.1 (signalling light COP9 signalosome): **P25** (VIT_202s0025g01170.1); MAPK belonging to Bincode 30.6 (signaling MAP kinases): **Q1** (VIT_200s0567g00010.1); Transcription Factors belonging to Bincode 27.3.25 (RNA regulation of transcription MYB domain transcription factor family): **R1** (VIT_214s0066g01090.1), **R2** (VIT_214s0108g00830.1) and **R3** (VIT_215s0046g00170.1); Heat Shot Proteins belonging to Bincode 20.2.1 (stress abiotic heat): **S1** (VIT_206s0004g04470.1), **S2** (VIT_215s0046g01560.1) and **S3** (VIT_219s0015g01370.1); Secondary metabolites belonging to Bincode 16.1 (secondary metabolism isoprenoids): **T1** (VIT_209s0054g01220.1), to Bincode 16.1.5. (secondary metabolism isoprenoids terpenoids): **T2** (VIT_209s0054g01220.1), **T3** (VIT_212s0134g00030.1) and **T4** (VIT_218s0001g04780.1), to Bincode 16.2.1.1 (secondary metabolism phenylpropanoids lignin biosynthesis PAL): **T5** (VIT_213s0019g04460.1), to Bincode 16.2.1.6 (secondary metabolism phenylpropanoids lignin biosynthesis CCoAOMT): **T6** (VIT_201s0010g03490.1) and **T7** (VIT_201s0010g03510.1), to Bincode 16.2.1.9 (secondary metabolism phenylpropanoids lignin biosynthesis COMT): **T8** (VIT_215s0048g02490.1), to Bincode 16.8.1.1 (secondary metabolism flavonoids anthocyanins leucocyanidin dioxygenase): **T9** (VIT_202s0025g04720.1), to Bincode 16.8.2 (secondary metabolism flavonoids chalcones): **T10** (VIT_210s0042g00840.1) and **T11** (VIT_210s0042g00920.1), to Bincode 16.8.2.1 (secondary metabolism flavonoids chalcones naringenin-chalcone synthase): **T12** (VIT_205s0136g00260.1), **T13** (VIT_210s0042g00880.1) and **T14** (VIT_210s0042g00920.1), to Bincode 16.8.3.1 (secondary metabolism flavonoids dihydroflavonols dihydroflavonol 4-reductase): **T15** (VIT_218s0001g12800.1), to Bincode 16.8.3.2 (secondary metabolism flavonoids dihydroflavonols flavanone 3-hydroxylase): **T16** (VIT_204s0023g03370.1) and **T17** (VIT_218s0001g14310.1), to Bincode 16.8.3.3 (secondary metabolism flavonoids dihydroflavonols flavonoid 3"-monooxygenase): **T18** (VIT_206s0009g02805.1), **T19** (VIT_206s0009g02810.1), **T20** (VIT_206s0009g02830.1), **T21** (VIT_206s0009g02840.1), **T22** (VIT_206s0009g02846.1), **T23** (VIT_206s0009g02860.1), T**24** (VIT_206s0009g02873.1), T**25** (VIT_206s0009g02880.1), T**26** (VIT_206s0009g02920.1), **T27** (VIT_206s0009g02970.1), **T28** (VIT_206s0009g03010.1) and to Bincode 16.8.4.3 (secondary metabolism flavonoids flavonols flavonol-3-O-rhamnosyltransferase): T29 (VIT_216s0039g02230.1).

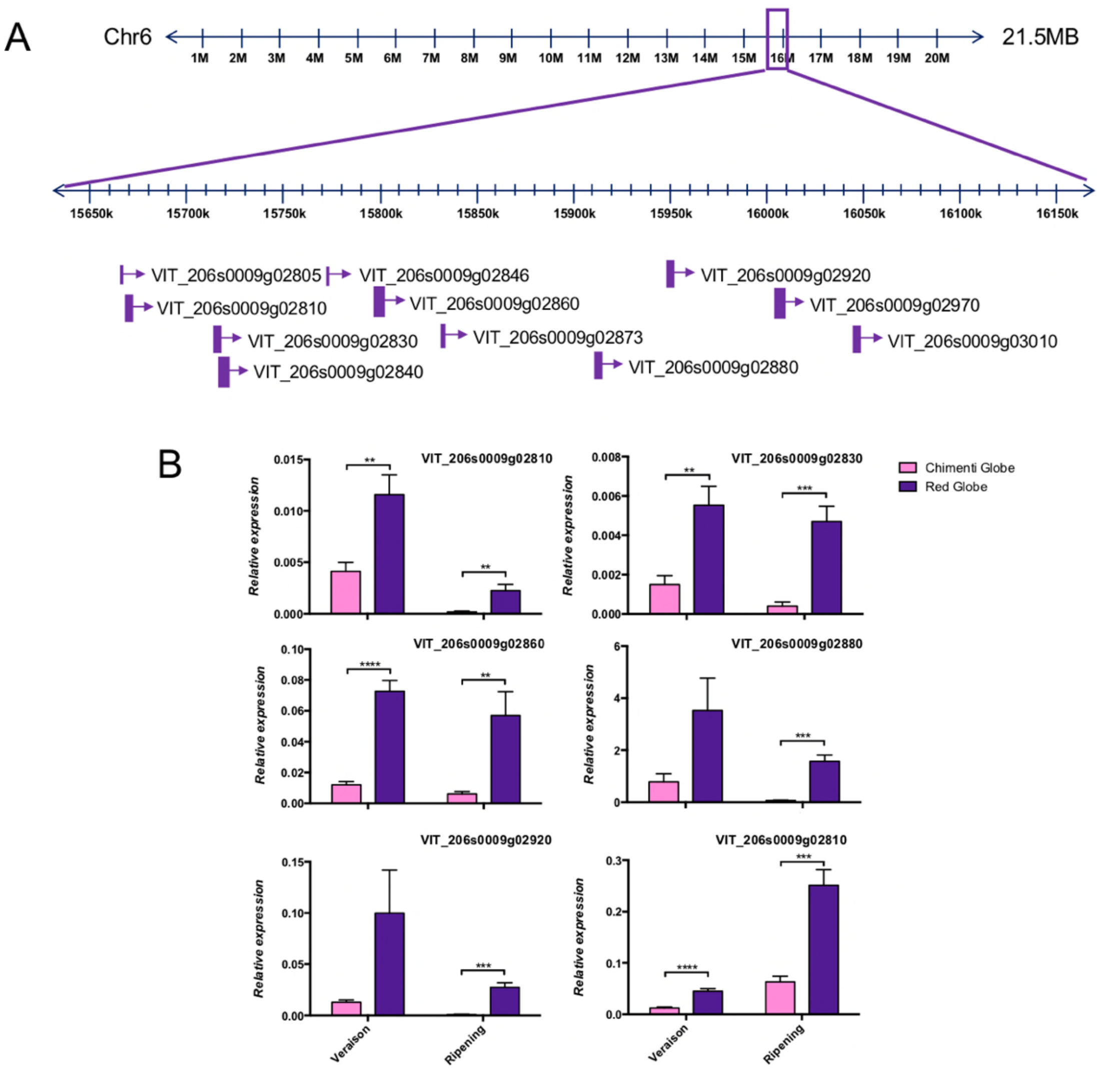


**FIGURE S6: Real-time quantitative PCR (qRT-PCR) validation of 6 F3'5'-H gene copies identified as repressed in the RNA-seq analysis of cv. 'Chimenti Globe'. (A) Genomic localization of 11 of the 15 gene copies of flavonoid 3'5'-hydroxylases (F3'5'-H) on chromosome 6 of *Vitis vinifera.*** Chromosome 6, with an approximate size of 21.5MB, possess fifteen F3'5'-H gene copies residing in a tandem array within a 650-kb region (Falginella et al., 2010). **(B)** Gene expression quantifications. Samples were collected at veraison and ripening stages in 2013 season. The gene copies of *F3'5-H* correspond to VIT_206s0009g02810, VIT_206s0009g02830, VIT_206s0009g02860, VIT_206s0009g02880, VIT_206s0009g02920 and VIT_206s0009g03010. Relative gene expressions are the mean of six biological replicates, with error bars representing the standard error of the mean. Asterisks indicate the result of unpaired T-tests between CG and RG. **, $P < 0.0085$; ***, $P < 0.0003$; ****, $P < 0.0001$.

**TABLE S1:** Pool designation of CG and RG plants for RNA-seq analysis.

| SKIN SAMPLE ID | POOL NAME |
|---|---|
| Veraison Chimenti Globe 1 | |
| Veraison Chimenti Globe 2 | POOL-1A |
| Veraison Chimenti Globe 3 | |
| Veraison Chimenti Globe 4 | POOL-1B |
| Veraison Chimenti Globe 5 | |
| Veraison Chimenti Globe 6 | POOL-1C |
| Veraison Red Globe 1 | |
| Veraison Red Globe 2 | POOL-2A |
| Veraison Red Globe 3 | |
| Veraison Red Globe 4 | POOL-2B |
| Veraison Red Globe 5 | |
| Veraison Red Globe 6 | POOL-2C |
| Ripening Chimenti Globe 1 | |
| Ripening Chimenti Globe 2 | POOL-3A |
| Ripening Chimenti Globe 3 | |
| Ripening Chimenti Globe 4 | POOL-3B |
| Ripening Chimenti Globe 5 | |
| Ripening Chimenti Globe 6 | POOL-3C |
| Ripening Red Globe 1 | |
| Ripening Red Globe 2 | POOL-4A |
| Ripening Red Globe 3 | |
| Ripening Red Globe 4 | POOL-4B |
| Ripening Red Globe 5 | |
| Ripening Red Globe 6 | POOL-4C |

**TABLE S2:** Primers of each gene analyzed by qRT-PCR and PCR

| Name | ID | Fw sequence | Rv sequence | % efficiency | TM (ºC) | Amplicon length (bp) |
|---|---|---|---|---|---|---|
| F3'5'-H multicopies | VIT_206s0009g02805<br>VIT_206s0009g02810<br>VIT_206s0009g02840<br>VIT_206s0009g02860<br>VIT_206s0009g02873<br>VIT_206s0009g02880<br>VIT_206s0009g02920<br>VIT_206s0009g02970<br>VIT_206s0009g03010 | TGGATAAAGTGATTGGAAGGA | GGATGGGTGCTTCCGGAAGCTT | 99,1 | 58 | 104 |
| CYTB5 | VIT_218s0001g09400 | GGCTCCCCAAAAAGTTTTCA | TGCCATGAATGACGAACCAG | 106,9 | 55 | 192 |
| UFGT | VIT_216s0039g02230 | GCTTTGATCAAATTCTCTCACA | ACACTAAATCCACCAGGGTTTT | 96,5 | 53 | 138 |
| AOMT2 | VIT_201s0010g03490 | TTGATCCAGATAAAGAAGCGTA | CGGCAATGAGATCATTTAGAAC | 101,3 | 55 | 120 |
| AOMT1 | VIT_201s0010g03510 | | | | | |
| AM1 | VIT_216s0050g00900 | ATTCGGGTCTCCAATGAGCT | GTGAGAAACAGCCTCTTGCA | 102,8 | 55 | 180 |
| AM2 | VIT_216s0050g00910 | | | | | |
| GST13 | VIT_207s0104g01800 | AATTCCCATGTCCACATGCACC | GGGATTCTTGGCAAGAAAAGGA | 98,8 | 56 | 130 |
| F3-H | VIT_204s0023g03370 | AGGCAACCGTGTATCCTCTG | TTCTCCACGTCTTGCAACTG | 102,9 | 60 | 160 |
| CHS | VIT_205s0136g00260 | CCCATGTTCGAATTGGTTTC | TGTGCAATCCAGAAAATGGA | 103,6 | 60 | 206 |
| MYB24 | VIT_214s0066g01090 | ACCCTGCAATTCTCAGGATG | CAGGCCTCAGGTAGTTCAGC | 100,2 | 60 | 182 |
| COMT | VIT_215s0048g02480 | TACCCTGGCAAGGACCCCAGAT | AGGTGCTCAAAACCCTTGTAGG | 100,3 | 60 | 110 |
| COMT | VIT_215s0048g02490 | | | | | |
| ABC TRANSPORTER | VIT_214s0060g00720 | AATGTGAAGAGCGAGGTGCT | CGACTGCCTTTCGTTCTTTC | 102,3 | 55 | 228 |
| STS1 | VIT_210s0042g00840 | TGTGCTCTGAGATTACTGTTGT | CAATGGAGGTATCTGGATCCGA | 99,5 | 53 | 132 |

| | | | | | | |
|---|---|---|---|---|---|---|
| VvSRP60 | VIT_205s0077g02060 | ATCTACCTCAAGCTCCTAGTC | CAATCTTGTCCTCCTTTCCT | 103 | 55 | 165 |
| F3'5'-H | VIT_206s0009g02810 | AGCATGGTGGTGGCCTCAACTC | TAGCTTTCTCAGTAGCTTCCAC | 109,9 | 60 | 165 |
| F3'5'-H | VIT_206s0009g02830 | GACCGGTTATTAACAAAGATGA | TGTCTGTTCCAGGAGGAGTGCT | 103,5 | 60 | 137 |
| F3'5'-Hf | VIT_206s0009g02860 | CACTAAGCCATGGCCATAGATA | GCAACATGAGGCATGTTACCTA | 100,7 | 60 | 176 |
| F3'5'-Hi | VIT_206s0009g02880 | AGTCAAATGAGTTCAAGGACATGG | AGGAGCACACGGCGTCAGCCCA | 101,2 | 60 | 131 |
| F3'5'-H | VIT_206s0009g02920 | TTTGGATCCTTCAAATATTC | GAAAAGATGGAGATGGCTCT | 100 | 55 | 125 |
| F3'5'-H | VIT_206s0009g03010 | TAACGGTATAAGATGTATTAGC | GAAAAATAGAAACTATAACCAA | 104,6 | 60 | 85 |
| VvMYBA1 white allele | VIT_202s0033g00410 | | | - | 65 | |
| VvMYBA1 red allele | VIT_202s0033g00410 | | | - | 65 | |
| VvACTIN | VIT_204s0044g00580 | ACCAGAATCCAGCACAATC | ATGGCCGATACTGAAGATAT | - | 55 | 444 |

**TABLE S3: Average content of different types of anthocyanins in Chimenti Globe and Red Globe berry skin (mg $g^{-1}$ DW).**

| | CHIMENTI GLOBE | | | | RED GLOBE | | | |
|---|---|---|---|---|---|---|---|---|
| | Veraison | | Ripening | | Veraison | | Ripening | |
| | ***2013*** | ***2014*** | ***2013*** | ***2014*** | ***2013*** | ***2014*** | ***2013*** | ***2014*** |
| Delphinidin | 0.00 ± 0.00[a] | 0.00 ± 0.00[a] | 0.00 ± 0.00[a] | 0.01 ± 0.00[a] | 0.07 ± 0.02[e] | 0.09 ± 0.09[b] | 0.52 ± 0.08[e] | 0.53 ± 0.03[c] |
| Petunidin | 0.00 ± 0.00[a] | 0.00 ± 0.00[a] | 0.01 ± 0.01[a] | 0.01 ± 0.00[a] | 0.06 ± 0.02[e] | 0.09 ± 0.09[a] | 0.60 ± 0.11[e] | 0.48 ± 0.23[c] |
| Malvidin | 0.00 ± 0.00[a] | 0.00 ± 0.00[a] | 0.01 ± 0.01[a] | 0.03 ± 0.01[a] | 0.21 ± 0.06[e] | 0.32 ± 0.37[a] | 2.07 ± 0.57[e] | 1.38 ± 0.64[c] |
| Cianidin | 0.11 ± 0.03[a] | 0.04 ± 0.02[a] | 0.69 ± 0.49[a] | 0.65 ± 0.14[a] | 0.39 ± 0.19[c] | 0.65 ± 0.26[d] | 2.17 ± 0.24[d] | 2.43 ± 0.65[d] |
| Peonidin | 0.16 ± 0.04[a] | 0.06 ± 0.02[a] | 1.21 ± 0.40[a] | 1.13 ± 0.32[a] | 0.75 ± 0.33[c] | 1.41 ± 0.86[c] | 9.04 ± 1.22[e] | 7.90 ± 2.08[e] |
| **Total** | **0.27 ± 0.05[a]** | **0.11 ± 0.04[a]** | **1.93 ± 0.85[a]** | **1.84 ± 0.44[a]** | **1.49 ± 0.57[c]** | **2.57 ±1.57[c]** | **14.39 ± 1.21[e]** | **12.71 ± 3.48[d]** |

The letters represent the result of an unpaired T-test between Chimenti Globe and Red Globe in each developmental stage during 2013 and 2014 season: [a] represent no significant differences; [b] represent *, $P = 0.0497$; [c] represent **, $P < 0.0093$; [d] represent ***, $P < 0.0009$; [e] represent ****, $P < 0.0001$.

**TABLE S4: Cleaning raw reads using Trimmomatic**

| Sample ID | Input total reads pairs | Both Kept | Forward Only Kept | Reverse Only Kept | Dropped |
|---|---|---|---|---|---|
| Pool-1A | 4852418 | 4575605 | 178246 | 60885 | 37682 |
| | (100.00%) | (94.30%) | (3.67%) | (1.25%) | (0.78%) |
| Pool-1B | 5173857 | 4871122 | 201139 | 60598 | 40998 |
| | (100.00%) | (94.15%) | (3.89%) | (1.17%) | (0.79%) |
| Pool-1C | 5681319 | 5350777 | 213111 | 72260 | 45171 |
| | (100.00%) | (94.18%) | (3.75%) | (1.27%) | (0.80%) |
| Pool-2A | 6553961 | 6169343 | 246950 | 85694 | 51974 |
| | (100.00%) | (94.13%) | (3.77%) | (1.31%) | (0.79%) |
| Pool-2B | 6570611 | 6180772 | 258552 | 79087 | 52200 |
| | (100.00%) | (94.07%) | (3.93%) | (1.20%) | (0.79%) |
| Pool-2C | 5311185 | 5010252 | 197744 | 64427 | 38762 |
| | (100.00%) | (94.33%) | (3.72%) | (1.21%) | (0.73%) |
| Pool-3A | 8183865 | 7690225 | 309652 | 111186 | 72802 |
| | (100.00%) | (93.97%) | (3.78%) | (1.36%) | (0.89%) |
| Pool-3B | 5982329 | 5624673 | 232378 | 76608 | 48670 |
| | (100.00%) | (94.02%) | (3.88%) | (1.28%) | (0.81%) |
| Pool-3C | 5694271 | 5360934 | 213180 | 74459 | 45698 |
| | (100.00%) | (94.15%) | (3.74%) | (1.31%) | (0.80%) |
| Pool-4A | 7331098 | 6895816 | 286456 | 90148 | 58678 |
| | (100.00%) | (94.06%) | (3.91%) | (1.23%) | (0.80%) |
| Pool-4B | 7604337 | 7147979 | 298053 | 95992 | 62313 |
| | (100.00%) | (94.00%) | (3.92%) | (1.26%) | (0.82%) |
| Pool-4C | 6606565 | 6229341 | 241663 | 84617 | 50944 |
| | (100.00%) | (94.29%) | (3.66%) | (1.28%) | (0.77%) |

**TABLE S5: Alignment of RNA-seq reads to the Grapevine Reference Genome V2 using HISAT2**

| Sample ID | Total reads paired | Reads aligned 0 times | Reads aligned 1 time | Reads aligned >1 time |
|---|---|---|---|---|
| Pool-1A | 4575605 | 508068 | 3301327 | 766210 |
| | (100.00%) | (11.1%) | (72.15%) | (16.75%) |
| Pool-1B | 4871122 | 766210 | 3508905 | 797273 |
| | (100.00%) | (11.6%) | (72.03%) | (16.37%) |
| Pool-1C | 5350777 | 642477 | 3808111 | 900189 |
| | (100.00%) | (12.01%) | (71.07%) | (16.82%) |
| Pool-2A | 6169343 | 670998 | 4482441 | 1015904 |
| | (100.00%) | (10.88%) | (72.66%) | (16.47%) |
| Pool-2B | 6180772 | 654089 | 4493751 | 1032932 |
| | (100.00%) | (10.58%) | (72.71%) | (16.71%) |
| Pool-2C | 5010252 | 868444 | 3387979 | 753829 |
| | (100.00%) | (17.33%) | (67.62%) | (15.05%) |
| Pool-3A | 7690225 | 1588674 | 4964146 | 1137405 |
| | (100.00%) | (20.66%) | (64.55%) | (14.79%) |
| Pool-3B | 5624673 | 883057 | 3820294 | 921322 |
| | (100.00%) | (15.70%) | (67.92%) | (16.38%) |
| Pool-3C | 5360934 | 715581 | 3748929 | 896424 |
| | (100.00%) | (13.35%) | (69.93%) | (16.72%) |
| Pool-4A | 6895816 | 791137 | 4928442 | 1176237 |
| | (100.00%) | (11.47%) | (71.47%) | (17.06%) |
| Pool-4B | 7147979 | 767762 | 5160674 | 1219543 |
| | (100.00%) | (10.74%) | (72.20%) | (17.06%) |
| Pool-4C | 6229341 | 729131 | 4436080 | 1064130 |
| | (100.00%) | (11.7%) | (71.21%) | (17.08%) |

**TABLE S6: Gene mapping analysis from aligned reads to Grapevine Reference Genome V2**

| Status | Pool-1A | Pool-1B | Pool-1C | Pool-2A | Pool-2B | Pool-2C | Pool-3A | Pool-3B | Pool-3C | Pool-4A | Pool-4B | Pool-4C |
|---|---|---|---|---|---|---|---|---|---|---|---|---|
| **Assigned** | 4166463 | 4429083 | 4850826 | 5613766 | 5621724 | 4087521 | 6382908 | 4990397 | 4860962 | 6323211 | 6573172 | 5691548 |
| **Unassigned_Ambiguity** | 0 | 0 | 0 | 0 | 0 | 0 | 0 | 0 | 0 | 0 | 0 | 0 |
| **Unassigned_MultiMapping** | 310015 | 321664 | 355914 | 403352 | 433391 | 299855 | 316418 | 363941 | 369011 | 545779 | 528626 | 499145 |
| **Unassigned_NoFeatures** | 129072 | 141636 | 140843 | 185174 | 179281 | 379406 | 408854 | 121440 | 92241 | 131729 | 139122 | 125051 |
| **Unassigned_Unmapped** | 154182 | 170911 | 214395 | 205819 | 204309 | 420061 | 770711 | 369383 | 260927 | 230418 | 225009 | 218959 |
| **Unassigned_MappingQuality** | 0 | 0 | 0 | 0 | 0 | 0 | 0 | 0 | 0 | 0 | 0 | 0 |
| **Unassigned_FragmentLength** | 0 | 0 | 0 | 0 | 0 | 0 | 0 | 0 | 0 | 0 | 0 | 0 |
| **Unassigned_Chimera** | 0 | 0 | 0 | 0 | 0 | 0 | 0 | 0 | 0 | 0 | 0 | 0 |
| **Unassigned_Secondary** | 0 | 0 | 0 | 0 | 0 | 0 | 0 | 0 | 0 | 0 | 0 | 0 |
| **Unassigned_Nonjunction** | 0 | 0 | 0 | 0 | 0 | 0 | 0 | 0 | 0 | 0 | 0 | 0 |
| **Unassigned_Duplicate** | 0 | 0 | 0 | 0 | 0 | 0 | 0 | 0 | 0 | 0 | 0 | 0 |

**TABLE S7: Reads from RNA-seq results for the 11 *F3'5-H* gene copies.**

| *Flavonoid 3'5' hydroxylase* | CG Veraison | RG Veraison | CG Ripening | RG Ripening |
|---|---|---|---|---|
| VITXXXX 2805 | 2 | 50 | 0 | 38 |
| VITXXXX 2810 | 15 | 91 | 0 | 65 |
| VITXXXX 2830 | 32 | 214 | 1 | 150 |
| VITXXXX 2840 | 18 | 186 | 1 | 135 |
| VITXXXX 2846 | 11 | 59 | 0 | 38 |
| VITXXXX 2860 | 9 | 68 | 15 | 153 |
| VITXXXX 2873 | 4 | 38 | 2 | 70 |
| VITXXXX 2880 | 2 | 50 | 1 | 38 |
| VITXXXX 2920 | 20 | 154 | 2 | 104 |
| VITXXXX 2970 | 12 | 106 | 1 | 81 |

**TABLE S8: Molecular marker analysis.**

| | | | SC8_010 | | SC8_026 | | VVNTM1 | | VVNTM2 | | VvMybA2 | | VvMybA1 | | VVNTM3 | | VVNTM5 | | VVNTM6 | | VVNTM4 | | VVIU20 | | VMC7G3 | |
|---|---|---|---|---|---|---|---|---|---|---|---|---|---|---|---|---|---|---|---|---|---|---|---|---|---|---|
| | Sample | Layer | 1 | 2 | 1 | 2 | 1 | 2 | 1 | 2 | 1 | 2 | 1 | 2 | 1 | 2 | 1 | 2 | 1 | 2 | 1 | 2 | 1 | 2 | 1 | 2 |
| Chimenti Globe | skin1 | L1+L2 | 121 | 125 | 239 | 254 | 165 | 171 | 376 | 389 | T | T | Gret | Non-Gret | 272 | 274 | 289 | 311 | 151 | 153 | 208 | 210 | 363 | 363 | 116 | 126 |
| | skin2 | | 121 | 125 | 239 | 254 | 165 | 171 | 376 | 389 | T | T | Gret | Non-Gret | 272 | 274 | 289 | 311 | 151 | 153 | 208 | 210 | 363 | 363 | 116 | 126 |
| | flesh4 | L2 | 121 | 125 | 239 | 254 | 165 | 171 | 389 | 389 | T | T | Gret | Non-Gret | 272 | 274 | 289 | 311 | 151 | 153 | 208 | 210 | 363 | 363 | 116 | 126 |
| | flesh5 | | 121 | 125 | 239 | 254 | 165 | 171 | 389 | 389 | T | T | Gret | Non-Gret | 272 | 274 | 289 | 311 | 151 | 153 | 208 | 210 | 363 | 363 | 116 | 126 |
| | flesh6 | | 121 | 125 | 239 | 254 | 165 | 171 | 389 | 389 | T | T | Gret | Non-Gret | 272 | 274 | 289 | 311 | 151 | 153 | 208 | 210 | 363 | 363 | 116 | 126 |
| | leaf7 | L1+L2 | 121 | 125 | 239 | 254 | 165 | 171 | 376 | 389 | T | T | Gret | Non-Gret | 272 | 274 | 289 | 311 | 151 | 153 | 208 | 210 | 363 | 363 | 116 | 126 |
| | leaf8 | | 121 | 125 | 239 | 254 | 165 | 171 | 376 | 389 | T | T | Gret | Non-Gret | 272 | 274 | 289 | 311 | 151 | 153 | 208 | 210 | 363 | 363 | 116 | 126 |
| Red Globe | skin9 | L1+L2 | 121 | 125 | 239 | 254 | 165 | 171 | 376 | 389 | T | T | Gret | Non-Gret | 272 | 274 | 289 | 311 | 151 | 153 | 208 | 210 | 363 | 363 | 116 | 126 |
| | skin10 | | 121 | 125 | 239 | 254 | 165 | 171 | 376 | 389 | T | T | Gret | Non-Gret | 272 | 274 | 289 | 311 | 151 | 153 | 208 | 210 | 363 | 363 | 116 | 126 |
| | skin11 | | 121 | 125 | 239 | 254 | 165 | 171 | 376 | 389 | T | T | Gret | Non-Gret | 272 | 274 | 289 | 311 | 151 | 153 | 208 | 210 | 363 | 363 | 116 | 126 |
| | flesh12 | L2 | 121 | 125 | 239 | 254 | 165 | 171 | 376 | 389 | T | T | Gret | Non-Gret | 272 | 274 | 289 | 311 | 151 | 153 | 208 | 210 | 363 | 363 | 116 | 126 |
| | flesh13 | | 121 | 125 | 239 | 254 | 165 | 171 | 376 | 389 | T | T | Gret | Non-Gret | 272 | 274 | 289 | 311 | 151 | 153 | 208 | 210 | 363 | 363 | 116 | 126 |
| | flesh14 | | 121 | 125 | 239 | 254 | 165 | 171 | 376 | 389 | T | T | Gret | Non-Gret | 272 | 274 | 289 | 311 | 151 | 153 | 208 | 210 | 363 | 363 | 116 | 126 |
| | leaf15 | L1+L2 | 121 | 125 | 239 | 256 | 165 | 171 | 376 | 389 | T | T | Gret | Non-Gret | 272 | 274 | 289 | 311 | 151 | 153 | 208 | 210 | 363 | 363 | 116 | 126 |
| | leaf16 | | 121 | 125 | 239 | 256 | 165 | 171 | 376 | 389 | T | T | Gret | Non-Gret | 272 | 274 | 289 | 311 | 151 | 153 | 208 | 210 | 363 | 363 | 116 | 126 |
| | leaf17 | | 121 | 125 | 239 | 254 | 165 | 171 | 376 | 389 | T | T | Gret | Non-Gret | 272 | 274 | 289 | 311 | 151 | 153 | 208 | 210 | 363 | 363 | 116 | 126 |

**TABLE S9: Reads observed for MYBA2 in the nucleotide position 141,805,14 in reference to the G/T substitution responsible for the gene's null function (Walker et al., 2016).**

| Pool | *T* | *G* | *C* | *A* |
|---|---|---|---|---|
| *1A* | *54* | | | |
| *1B* | *61* | | | |
| *1C* | *118* | | | |
| *2A* | *90* | | | |
| *2B* | *115* | | | |
| *2C* | *84* | *1* | | |
| *3A* | *0* | | | |
| *3B* | *21* | | | |
| *3C* | *39* | | | |
| *4A* | *66* | | | |
| *4B* | *127* | | | |
| *4C* | *68* | | | |
